## Supplementary Figures 1-26 for "The cellular state space of AML unveils novel *NPM1* subtypes with distinct clinical outcomes and immune evasion properties"

### SUPPLEMENTARY INFORMATION

**Supplementary figures**
**Supplementary references**

### Supplementary Figures

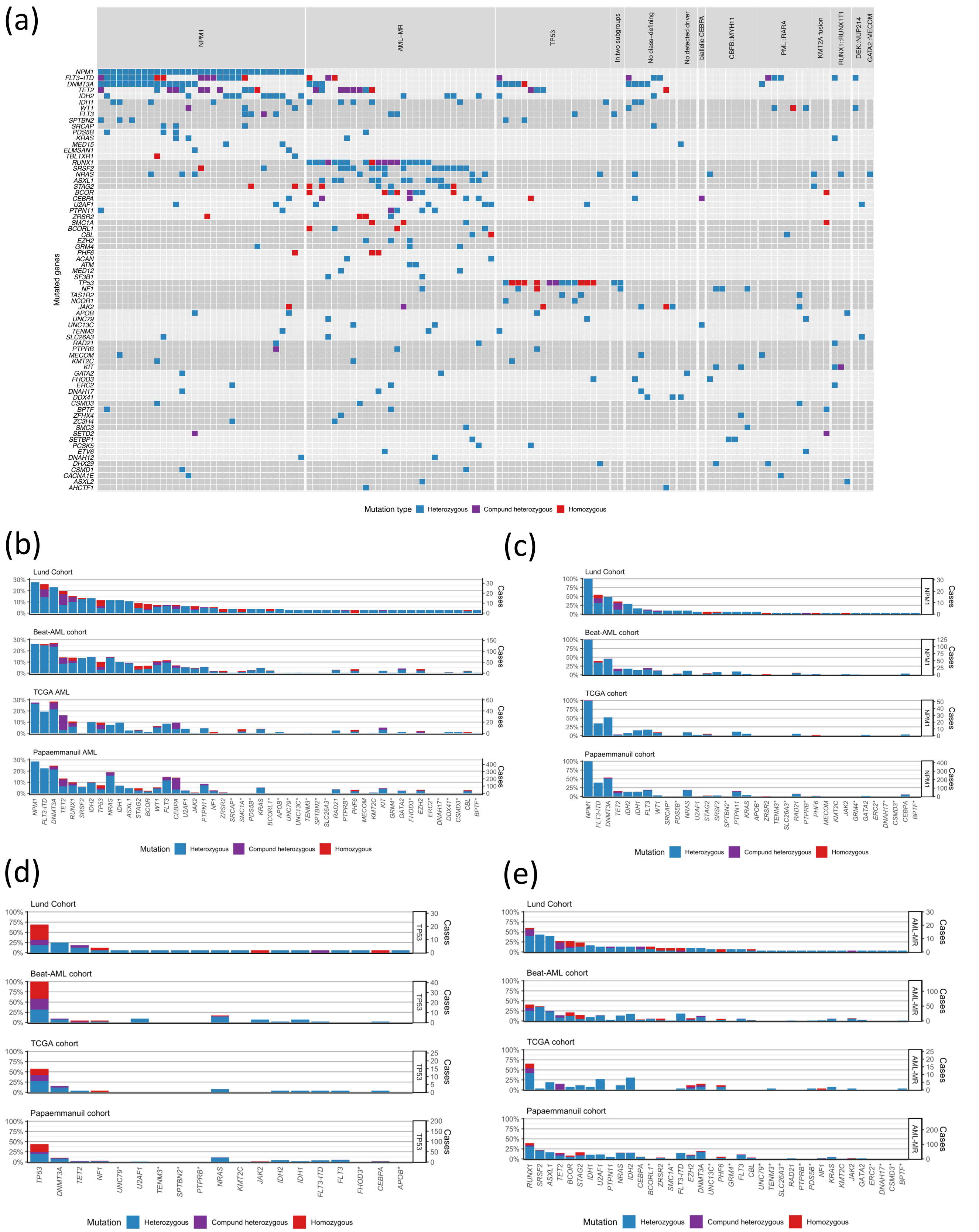


**Supplementary Fig. 1 | Somatic mutations in AML. (a)** Small nucleotide variants and insertion/deletions in the 70 most commonly mutated genes for the 120 AMLs included in the analyzed cohort. **(b)** Overall mutation frequencies compared with the AML cohorts from TCGA,[1] Beat-AML,[2] and Papaemmanuil et al[3]. **(c)** Mutation frequencies among NPM1-mutated cases compared with the AML cohorts from TCGA, Beat-AML, and Papaemmanuil et al. **(d)** Mutation frequencies among TP53-mutated cases compared with the AML cohorts from TCGA, Beat-AML, and Papaemmanuil et al. **(e)** Mutation frequencies among AML-MR cases compared with the AML cohorts from TCGA, Beat-AML, and Papaemmanuil et al.

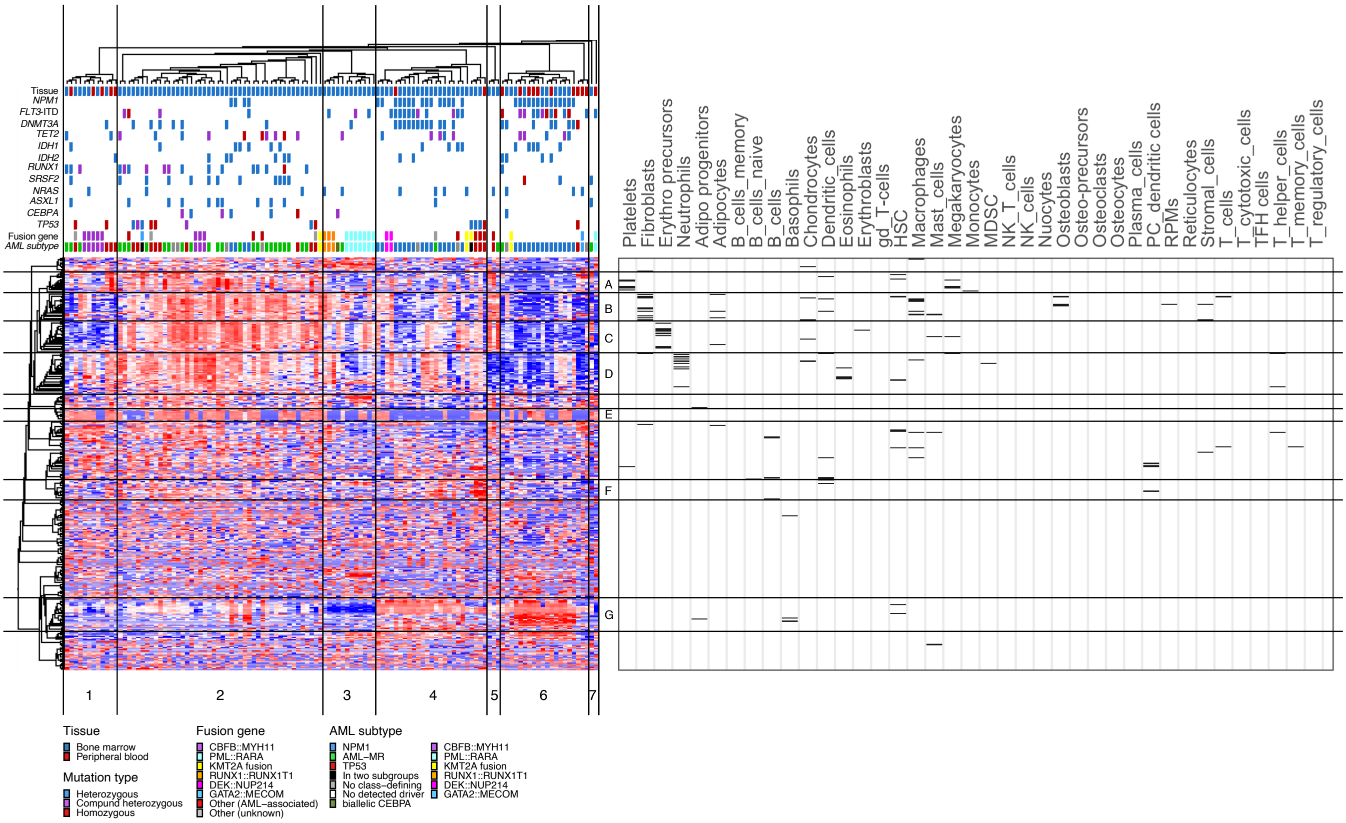


**Supplementary Fig. 2 | Cell type markers are enriched among the genes with most variable expression in AML.** Left side: Hierarchical clustering of the 120 AML samples based on gene expression of the 371 most variable genes (s/smax = 0.45). Distinct gene clusters are indicated by letters on the right side of the heatmap based on the gene dendogram. Right side: Genes within the heatmap that match cell type marker genes in PanglaoDB[4] for 39 distinct cell types present in blood, bone, connective tissue, or the immune system are indicated by black lines. In gene cluster A, 5/19 genes (26%) match marker genes for platelets. In cluster B, 9/25 genes (36%) match marker genes for fibroblasts. In gene cluster C, 8/26 genes (30%) match marker genes for erythroid-like and erythroid precursor cells. In gene cluster D, 8/33 genes (24%) match marker genes for neutrophils.

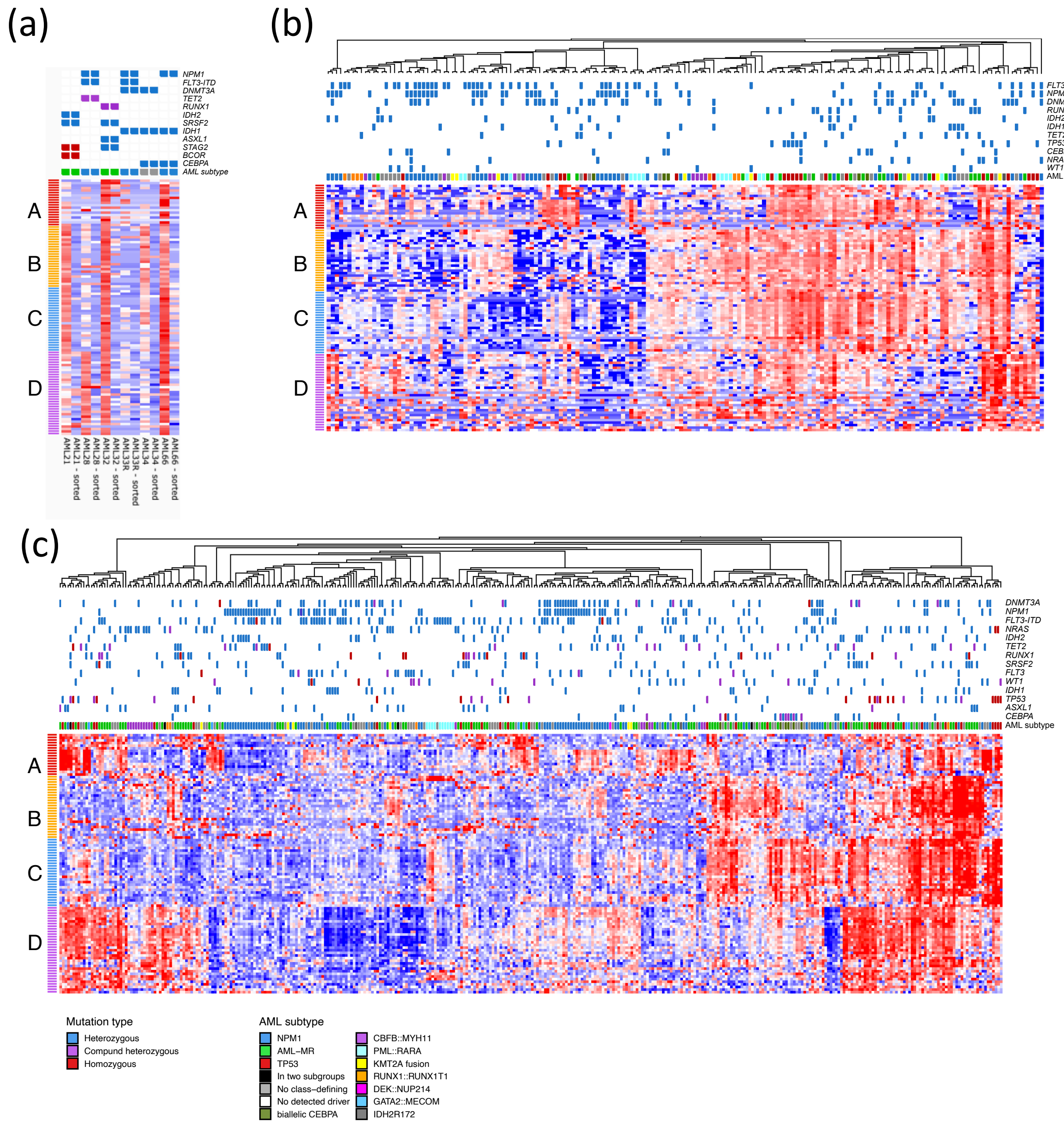


**Supplementary Fig. 3 | Cell type markers in sorted samples and external datasets. (a)** Heatmap illustrating the expression level of genes in the four gene clusters enriched for cell type markers (A,B,C, and D). Six bulk samples (AML21, AML28, AML32, AML33R, AML34, and AML66) and six samples sorted to contain only myeloid (CD33+/CD19-/CD3-) mononuclear cells (indicated with "- sorted") are included. **(b)** and **(c)** Heatmaps illustrating the expression level of genes in the four gene clusters enriched for cell type markers (A,B,C, and D) in the TCGA[1] **(b)** and Beat-AML[2] **(c)** cohorts.

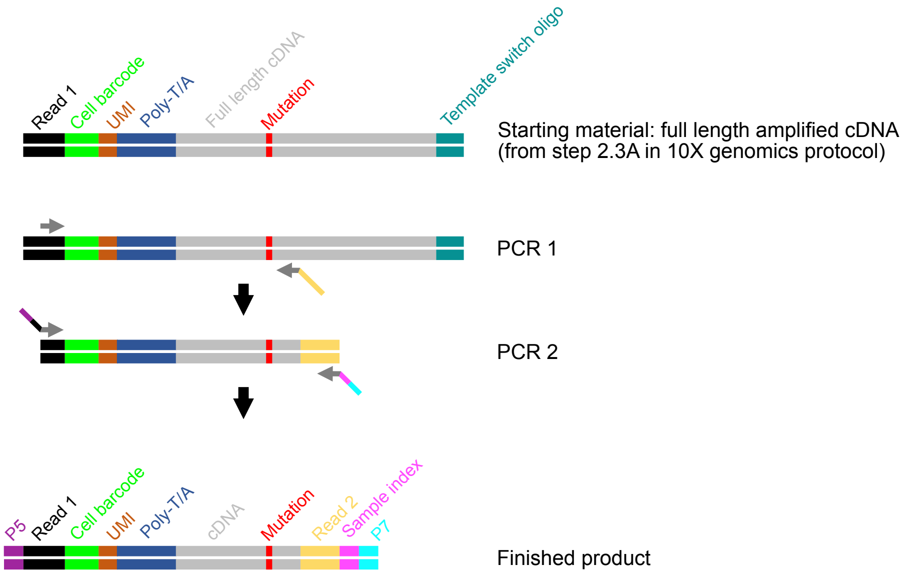


**Supplementary Fig. 4 | Overview of mutation calling PCR strategy.** Single cell mutation calling on scRNA-seq material was performed using a two-step PCR-amplification protocol. Full length amplified cDNA from an intermediate step in the 10X genomics chromium single cell 3´ v3 library preparation (step 2.3A) was used as PCR-template. In the first PCR, the right-hand primer was placed less than 100 bp from the targeted mutation. A 34-bp overhang (yellow) was added to the right-hand primer sequence. The left-hand primer for the first PCR was placed within the illumina Read1 sequence (black), thereby retaining the cell barcode and UMI information within the amplified material. In the second PCR, general primers binding to the left-hand Read1 sequence (black) and the right-hand Read2 sequence (yellow) were utilized, with overhang sequences adding sample index information (pink) and the P5 and P7 adapters (purple and turquoise) required for illumina sequencing. At sequencing, read 1 provides cell barcode and UMI information while read 2 provides mutation status for the targeted mutation.

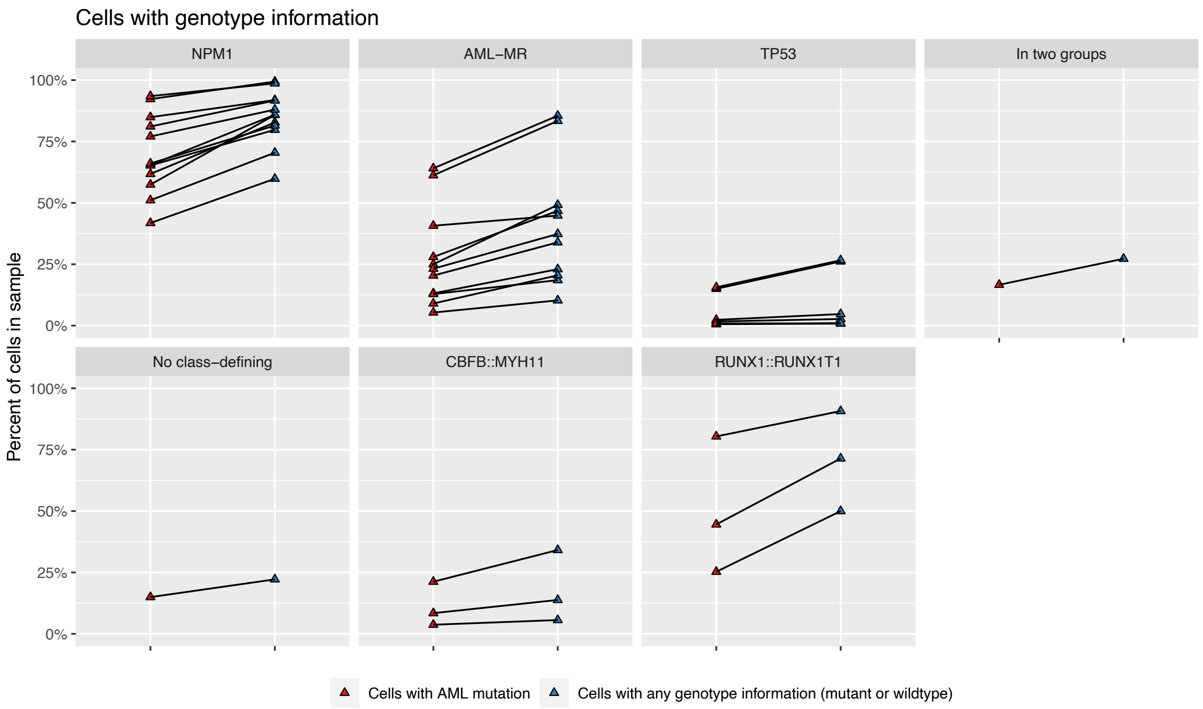


**Supplementary Fig. 5 | Proportion of cells with genotype information.** The proportion of cells with a detected AML mutation (red) and the proportion of cells with any genotype information (either mutant or wildtype) for the targeted positions (blue) illustrated for each sample. The samples are grouped by AML subtype.

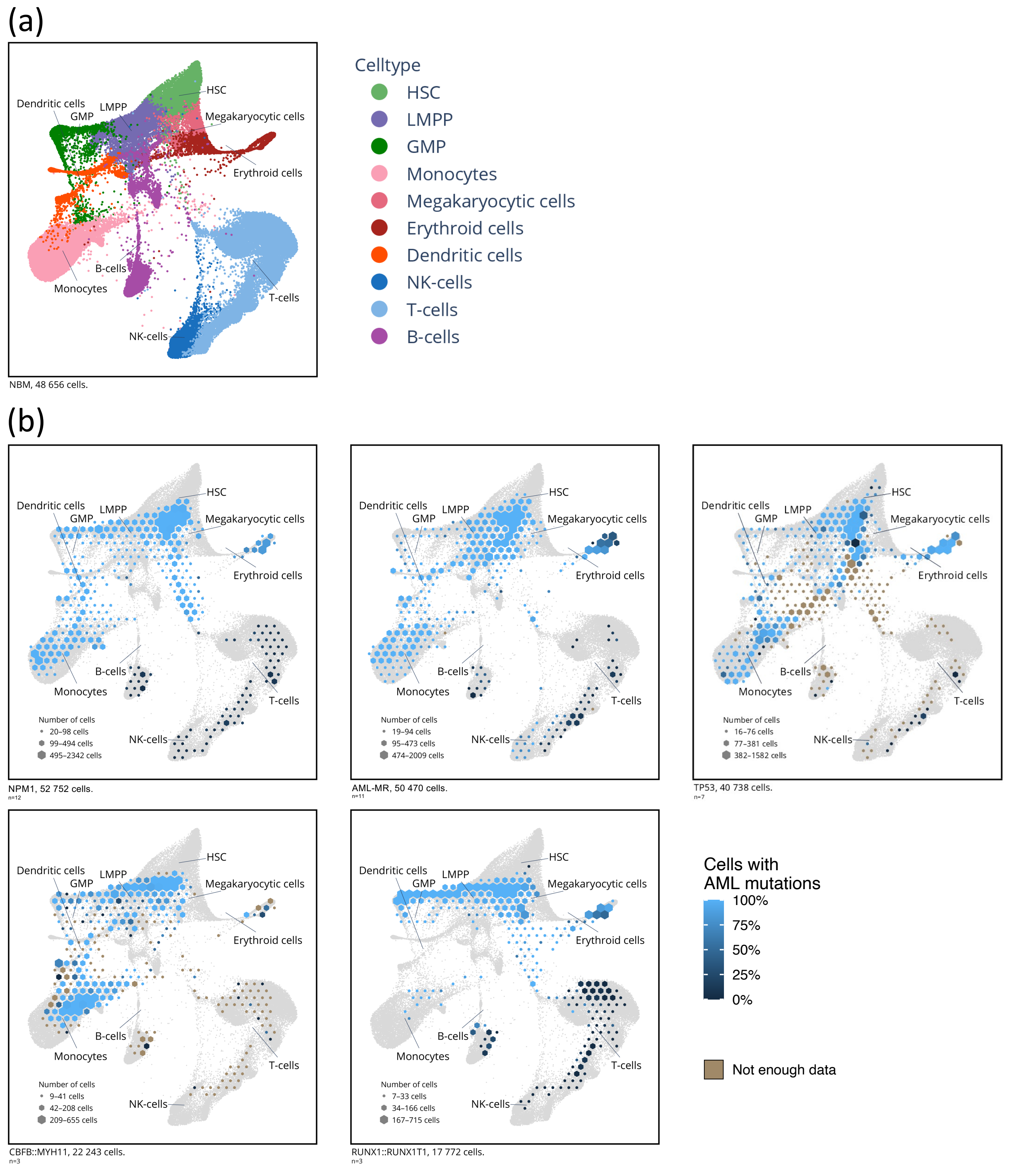


**Supplementary Fig. 6 | Single cell projection of AML subtypes onto NBM reference knn force plot. (a)** Knn force graph constructed from 42,370 NBM cells. **(b)** Projection of single cells onto the reference NBM knn force graph (indicated in gray), divided by AML subtype. The number of cells projected onto a region is indicated by the size of each pixel and the proportion of mutated cells in that pixel is indicated by color. Pixels with too few genotype reads are marked in brown (three or fewer reads). The genomic subtype of each sample is indicated below each plot.

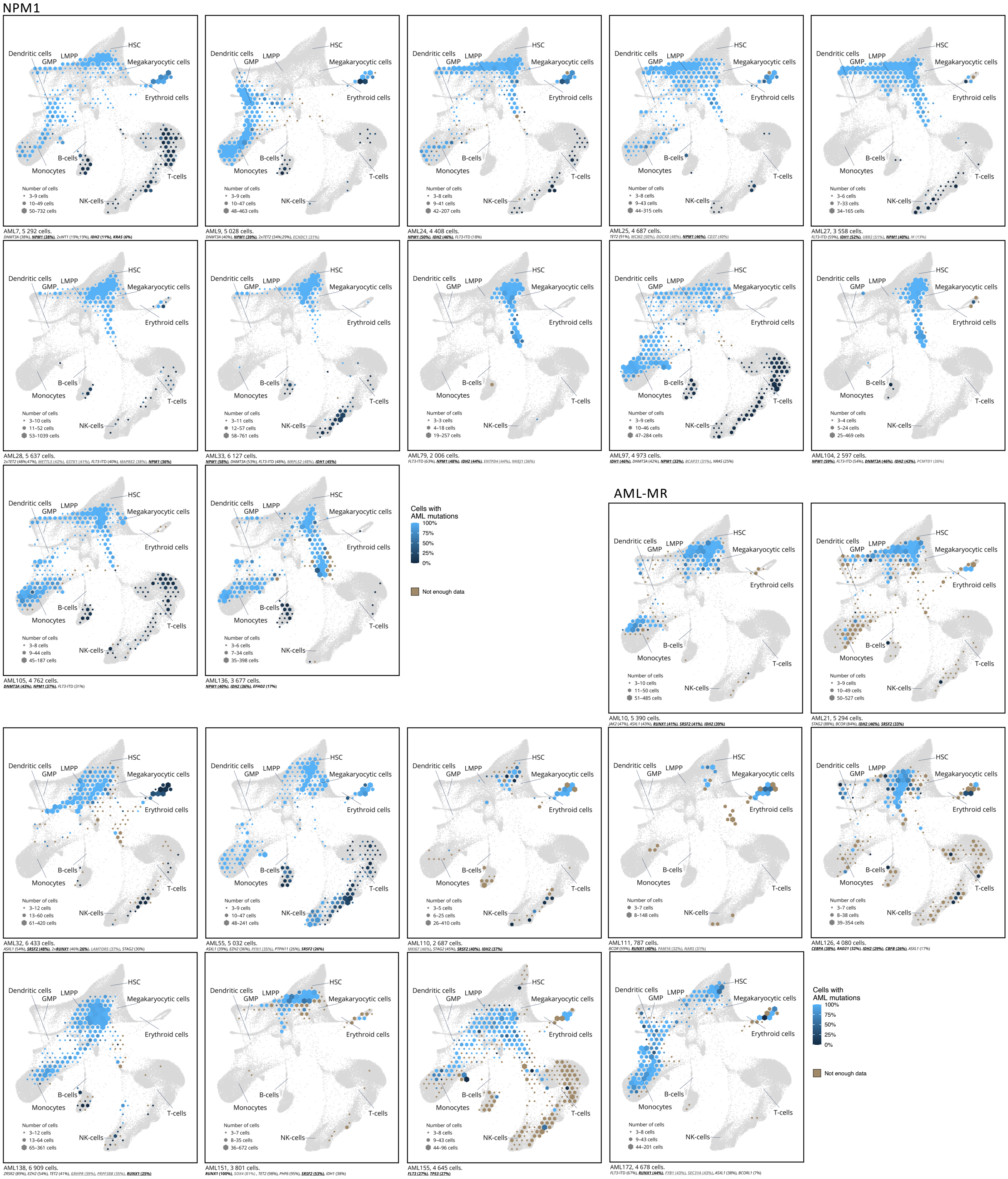


**Supplementary Fig. 7 | Single cell projection of individual AML samples from NPM1 and AML-MR subtypes onto NBM reference knn force plot.** Projection of single cells from AML samples onto the reference NBM knn force graph (indicated in gray), divided by sample. The number of cells projected onto a region is indicated by the size of each pixel and the proportion of mutated cells in that pixel is indicated by color. Pixels with too few genotype reads are marked in brown (three or fewer reads). Sample identifier and number of cells are indicated below each plot. A selection of genes mutated in each sample is presented below each plot, with the variant allele frequency indicated by WES denoted in parentheses. Genes targeted by scRNAmut-seq are indicated in bold. Genes with heterozygous mutations (used for inferring the proportion of mutated cells) are underlined. Genes with presumed passenger mutations are denoted in gray.

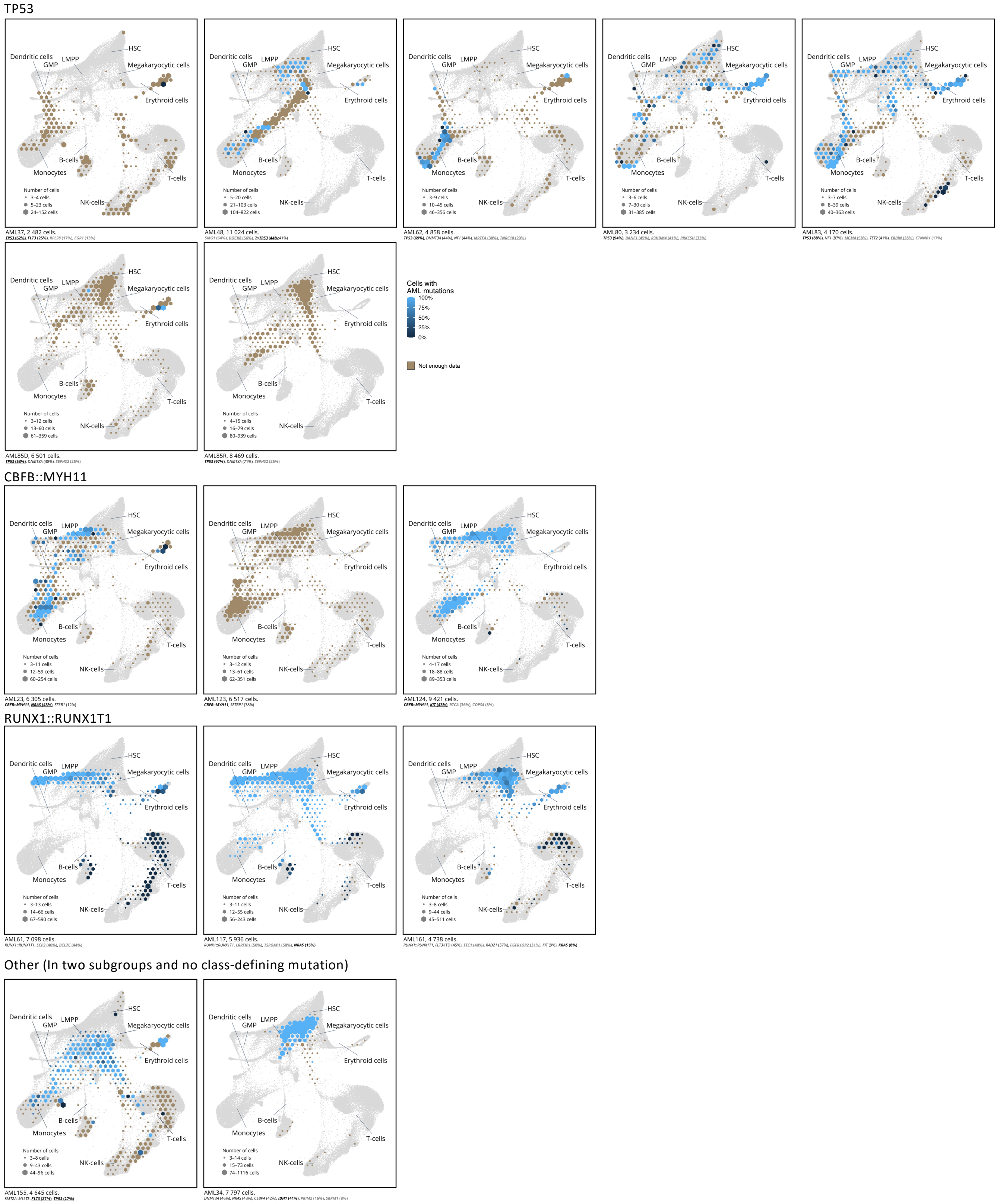


**Supplementary Fig. 8 | Single cell projection of individual AML samples from TP53, CBFB::MYH11, RUNX1::RUNX1T1 and other subtypes onto NBM reference knn force plot.** Projection of single cells from AML samples onto the reference NBM knn force graph (indicated in gray), divided by sample. The number of cells projected onto a region is indicated by the size of each pixel and the proportion of mutated cells in that pixel is indicated by color. Pixels with too few genotype reads are marked in brown (three or fewer reads). Sample identifier and number of cells are indicated below each plot. A selection of genes mutated in each sample is presented below each plot, with variant allele frequency from WES denoted in parentheses. Genes targeted by scRNAmut-seq are indicated in bold. Genes with heterozygous mutations (used for inferring the proportion of mutated cells) are underlined. Genes with presumed passenger mutations are denoted in gray.

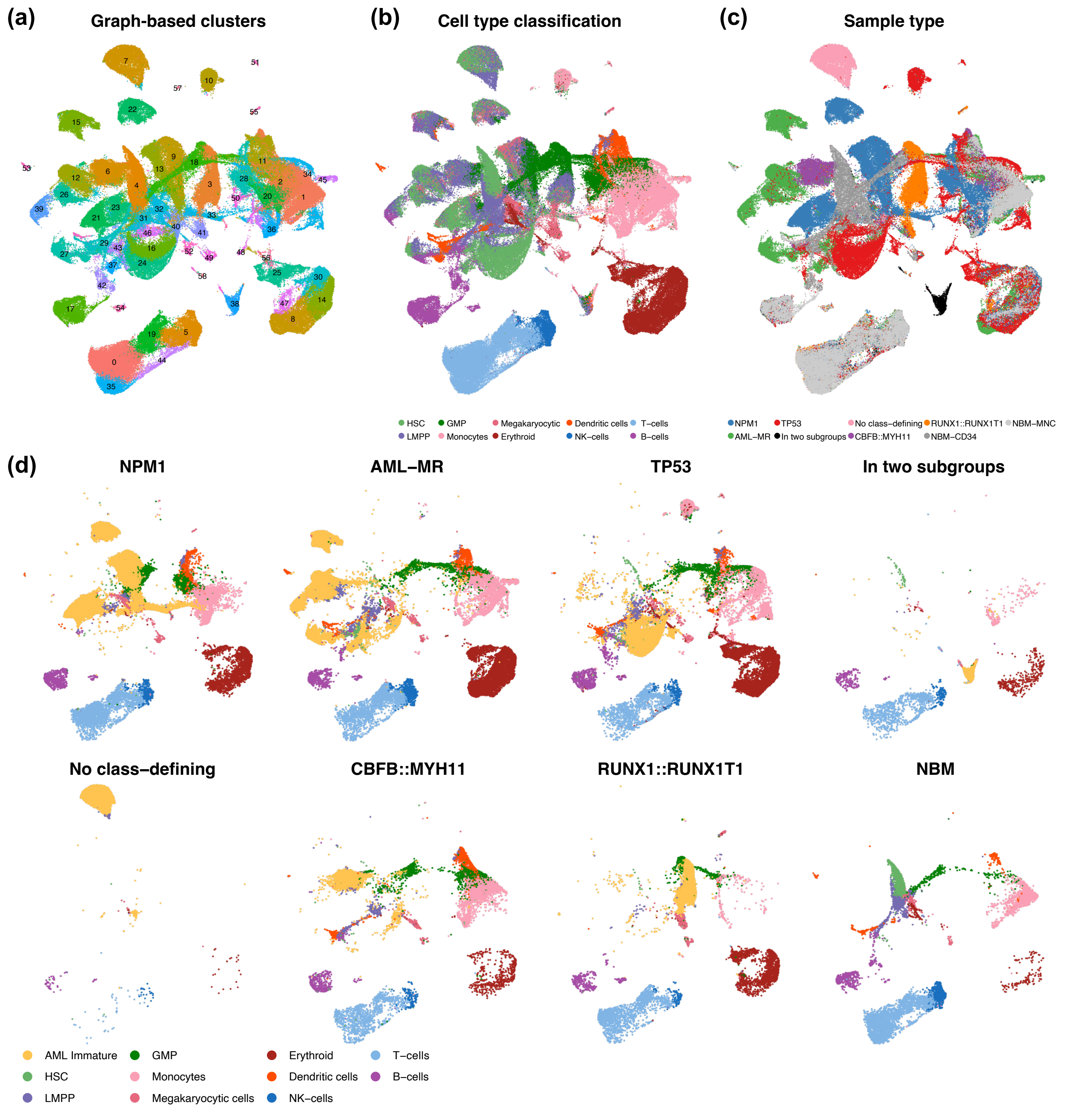


**Supplementary Fig. 9 | Classification of cell types and identification of AML immature cells visualized using UMAP. (a-c)** UMAP representation of 245,073 cells from 38 AML samples and 8 NBM samples. **(a)** Cells are colored based on graph-based clustering, which identified 58 distinct clusters of cells within the data. **(b)** Cells are colored by cell type based on classification using a reference dataset of NBM cells. **(c)** Cells are colored by AML subtype or NBM sample type. Notably, all AML samples were associated with a distinct cluster of cells outside the regions occupied by cells from NBM samples. The majority of cells in these AML-specific clusters were classified as HSC or LMPP based on the reference dataset. The cells in the AML-specific clusters were therefore reclassified as "AML immature". **(d)** UMAP representations of 245,073 cells from 38 AML samples and 8 NBM samples, separated into groups defined by AML subtype or NBM status. AML immature cells are indicated in yellow.

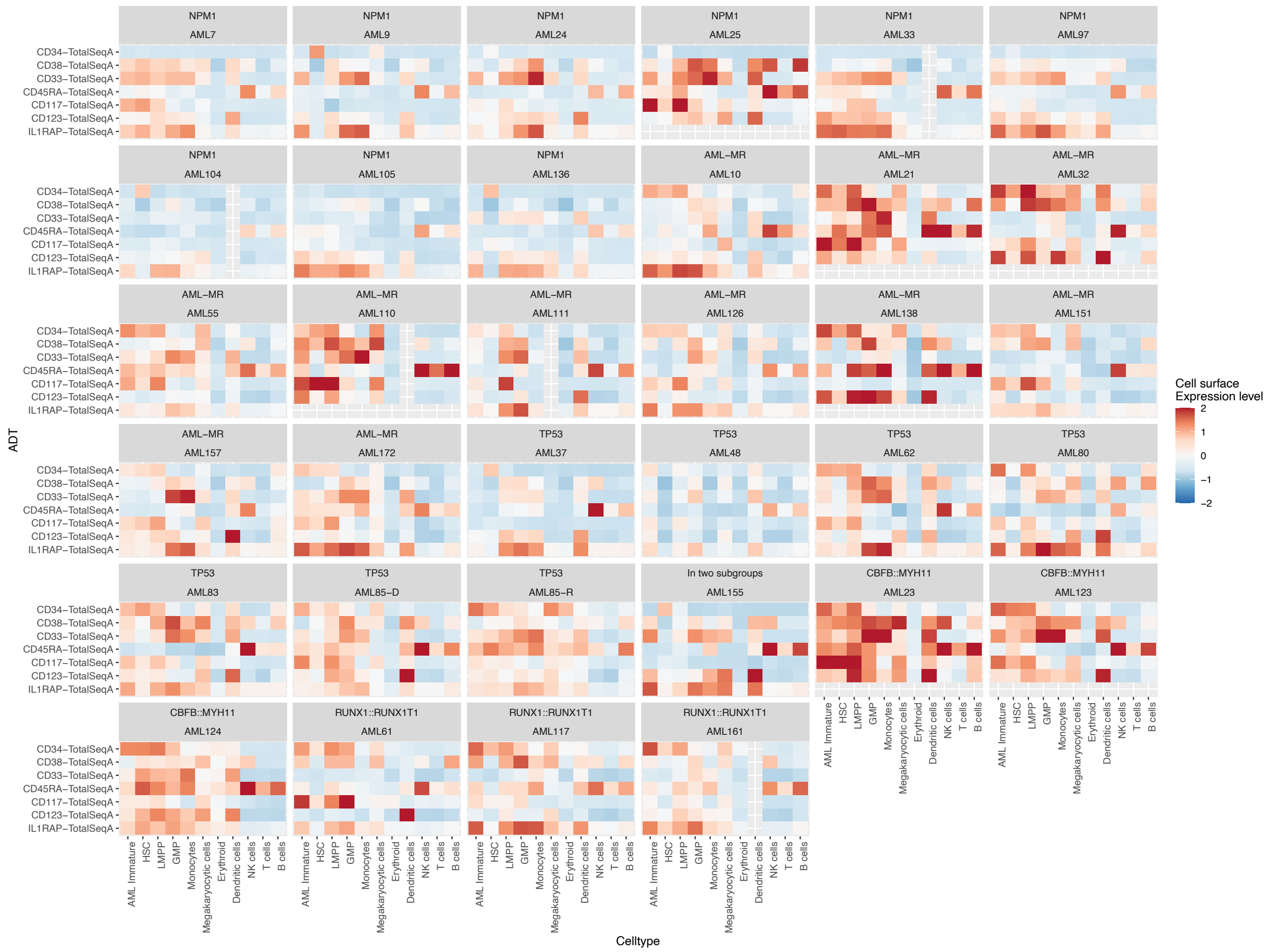


**Supplementary Fig. 10 | Immunophenotypic features of distinct cell types in 34 AML samples as determined by scADT-seq.** Heatmaps displaying the average cell surface expression of seven LSC markers for each cell type in 34 AML samples (scADT-seq data was not available from four AML samples; data for IL1RAP was only available from 27 AML samples). The AML immature cell type has an immunophenotype compatible with harboring LSC (i.e. $CD34^{+}CD38^{-/low}$ for CD34-positive AMLs and distinct expression profiles of CD33, CD45RA, CD117, CD123, and IL1RAP).

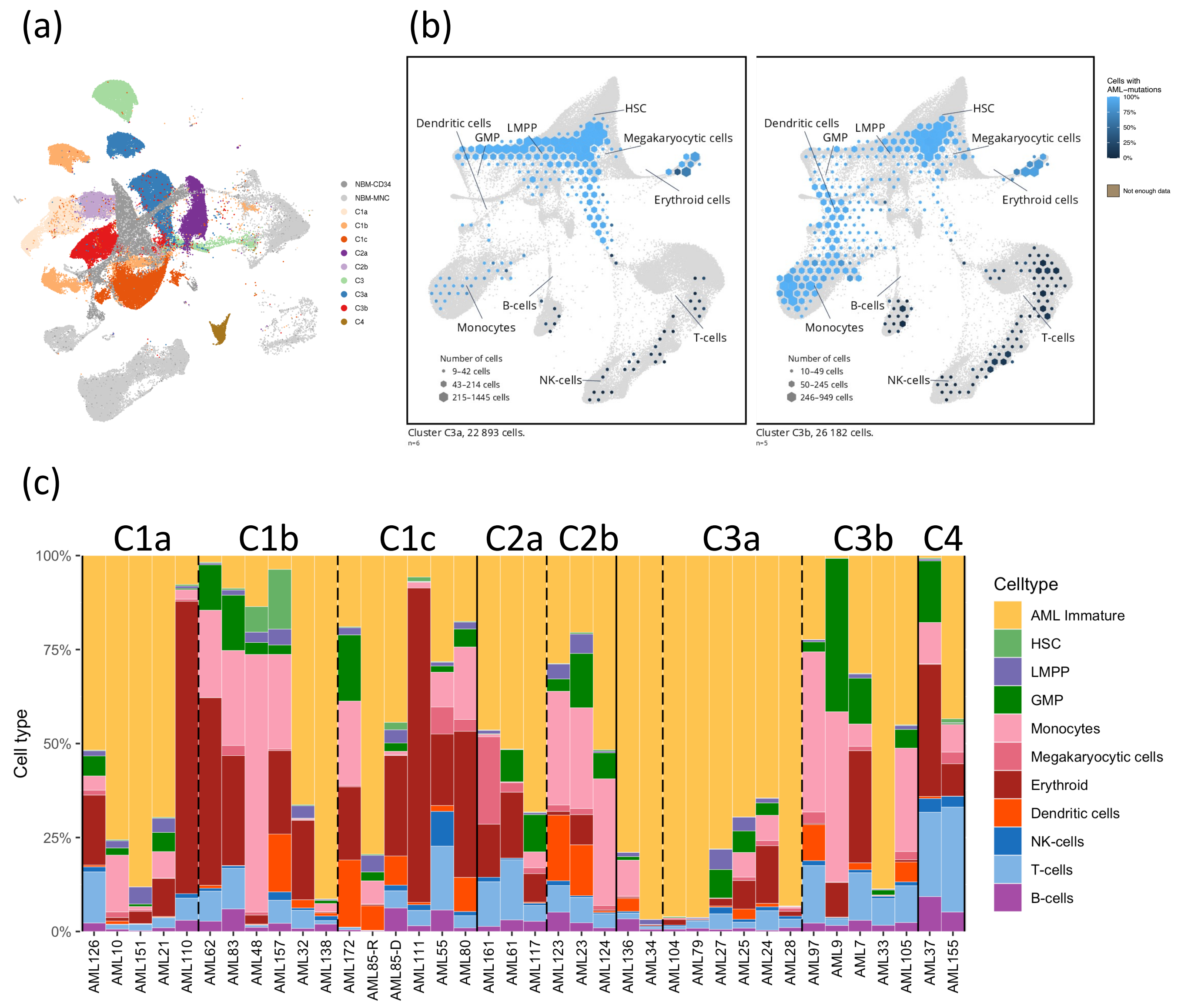


**Supplementary Fig. 11 | UMAP clustering and cell type distribution of AML immature clusters from hierarchical clustering analysis. (a)** UMAP representation of 245,073 cells from 38 AML samples and 8 NBM samples. Cells are colored by cluster identity of the AML sample based on hierarchical clustering on average gene expression of the AML immature cells from each AML sample (see Figure 3a). **(b)** Projection of single cells onto the reference NBM knn force graph (indicated in gray) from AML samples belonging to clusters C3a and C3b, separated by cluster identity. Samples in cluster C3a (left) contain a high proportion of immature cells whereas samples from cluster C3b (right) contain a higher proportion of T-cells and differentiated AML cells. **(c)** Cell type distribution in samples from each cluster defined by hierarchical clustering on average gene expression of the AML immature cells from each AML sample (Figure 3a). Samples from cluster C3a contain a high proportion of AML immature cells whereas samples from cluster C3b contain a high proportion of GMPs, monocytes, and T-cells.

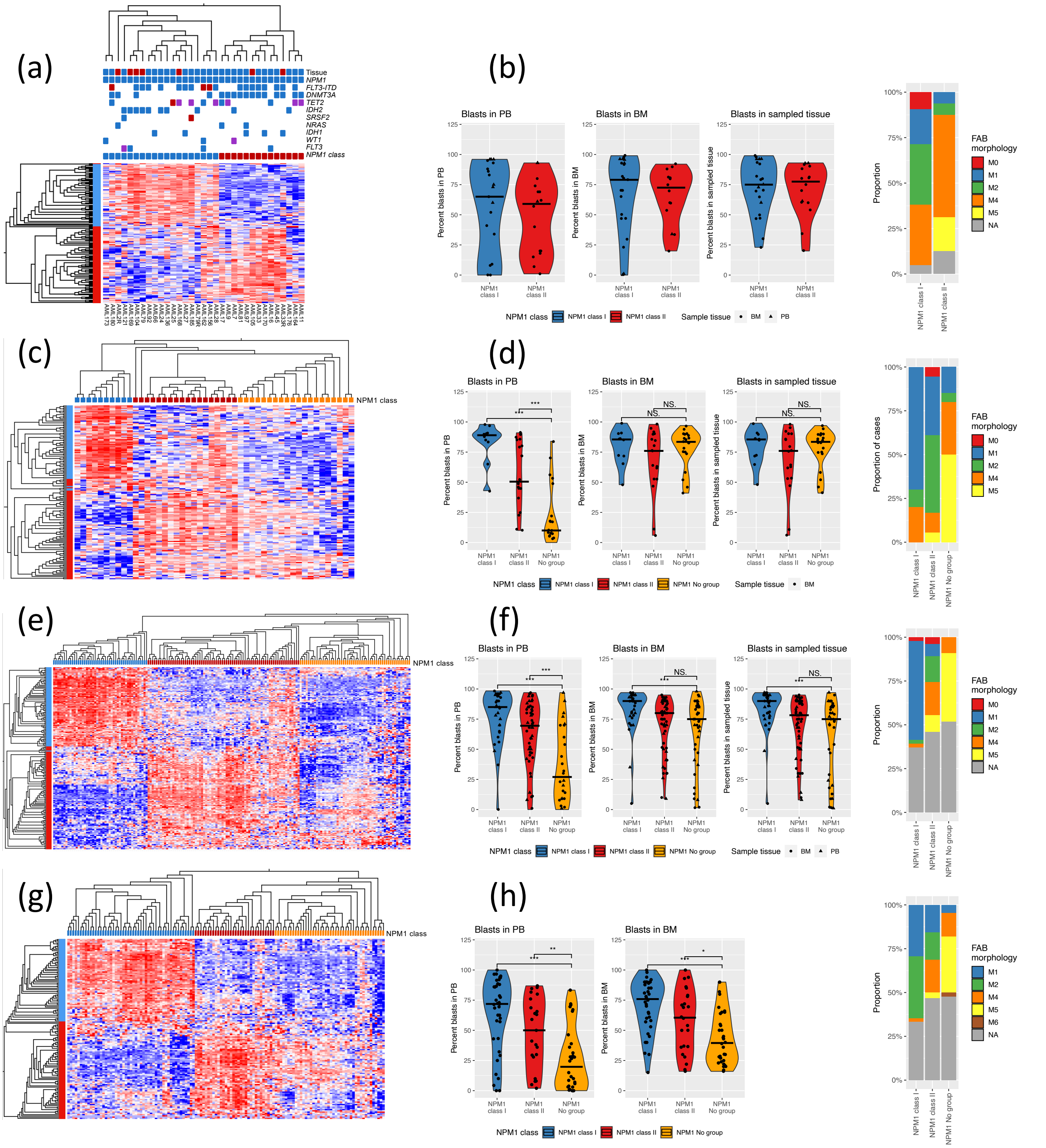


**Supplementary Fig. 12 | Identification of *NPM1*$^{\text{class I}}$ and *NPM1*$^{\text{class II}}$ in bulk RNA sequencing data. (a)** Hierarchical clustering of bulk gene expression data from 33 *NPM1*-mutated AML samples from the Lund dataset based on the expression of 180 genes differentially expressed between AML immature cells from *NPM1*$^{\text{class I}}$ (genes indicated in blue) and *NPM1*$^{\text{class II}}$ samples (genes indicated in red). Sample tissue and mutations in selected genes are indicated above each sample. In this dataset, 19 had an expression profile matching *NPM1*$^{\text{class I}}$ (samples indicated in blue) and 14 samples had an expression profile matching *NPM1*$^{\text{class II}}$ (samples indicated in red). **(b)** Percent blasts in PB, BM, and sample tissue (either PB or BM) for *NPM1*$^{\text{class I}}$ and *NPM1*$^{\text{class II}}$ cases from Lund (left side). FAB morphology for *NPM1*$^{\text{class I}}$ and *NPM1*$^{\text{class II}}$ cases from Lund (right side). **(c)** Hierarchical clustering of bulk gene expression data from 48 *NPM1*-mutated AML samples from the TCGA dataset[1] based on the expression of 180 genes differentially expressed between *NPM1*$^{\text{class I}}$ and *NPM1*$^{\text{class II}}$. **(d)** Percent blasts in PB, BM, and sample tissue (either PB or BM) for *NPM1*$^{\text{class I}}$, *NPM1*$^{\text{class II}}$, and unclassified cases from TCGA, with significant differences indicated by asterisks (Mann-Whitney U test; left side). FAB morphology for *NPM1*$^{\text{class I}}$, *NPM1*$^{\text{class II}}$, and unclassified cases from TCGA (right side). **(e)** Hierarchical clustering of bulk gene expression data from 174 *NPM1*-mutated AML samples from the Beat-AML 2.0 dataset[5] based on the expression of 180 genes differentially expressed between *NPM1*$^{\text{class I}}$ and *NPM1*$^{\text{class II}}$. **(f)** Percent blasts in PB, BM, and sample tissue (either PB or BM) for *NPM1*$^{\text{class I}}$, *NPM1*$^{\text{class II}}$, and unclassified cases from Beat-AML 2.0, with significant differences indicated by asterisks (Mann-Whitney U test; left side). FAB morphology for *NPM1*$^{\text{class I}}$, *NPM1*$^{\text{class II}}$, and unclassified cases from Beat-AML 2.0 (right side). **(g)** Hierarchical clustering of bulk gene expression data from 127 *NPM1*-mutated AML samples from the Clinseq dataset[6] based on the expression of 180 genes differentially expressed between *NPM1*$^{\text{class I}}$ and *NPM1*$^{\text{class II}}$. **(h)** Percent blasts in PB, BM, and sample tissue (either PB or BM) for *NPM1*$^{\text{class I}}$, *NPM1*$^{\text{class II}}$, and unclassified cases from Clinseq, with significant differences indicated by asterisks (Mann-Whitney U test; left side). FAB morphology for *NPM1*$^{\text{class I}}$, *NPM1*$^{\text{class II}}$, and unclassified cases from Clinseq (right side).

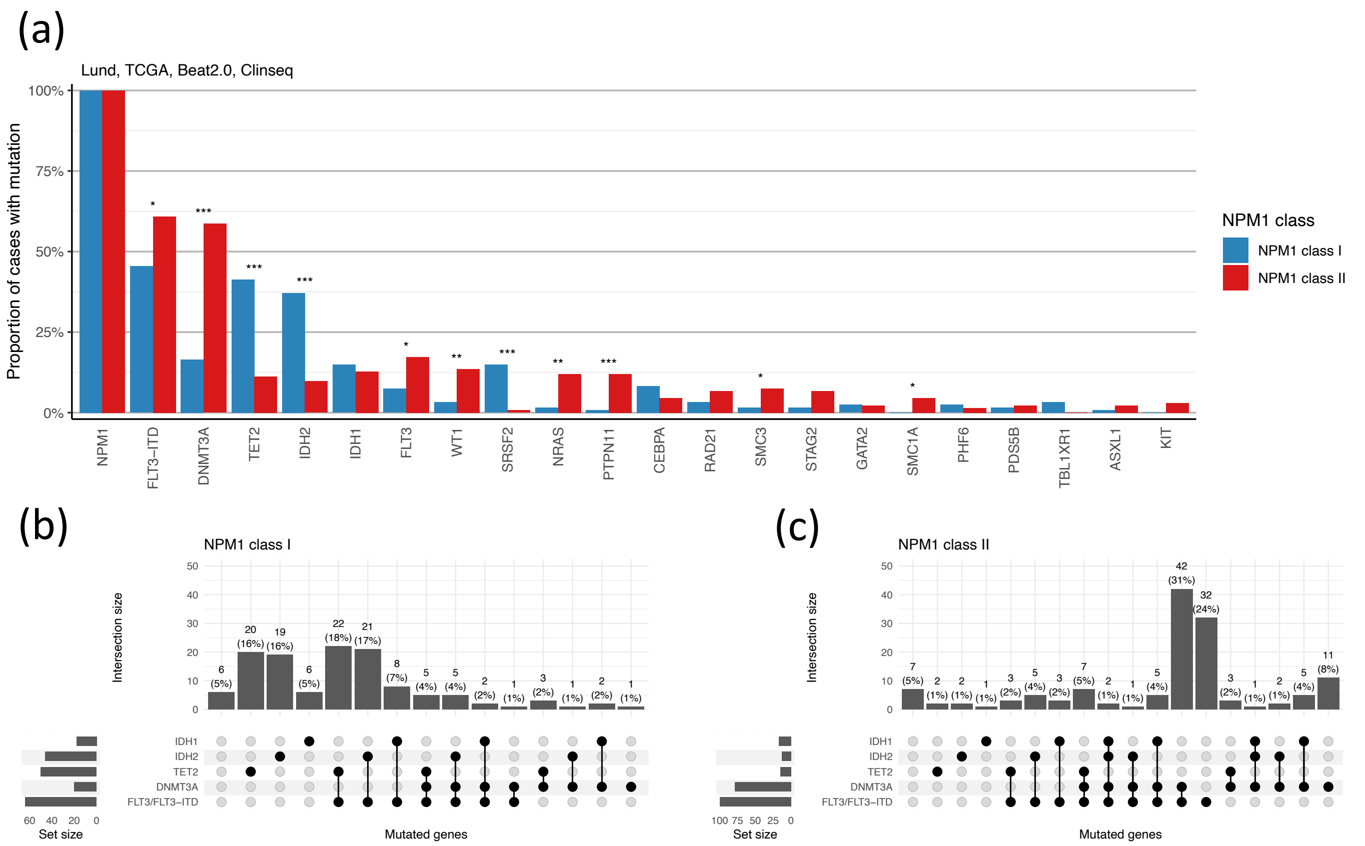


**Supplementary Fig. 13 | Mutational patterns of *NPM1*$^{\text{class I}}$ and *NPM1*$^{\text{class II}}$ subtypes. (a)** Proportion of cases with mutations in the most commonly mutated genes in the *NPM1*$^{\text{class I}}$ and *NPM1*$^{\text{class II}}$ subtypes**.** Significant differences between the groups are indicated with asterisks (*: p<0.05; **: p<0.01; ***: p<0.001; Fisher's exact test). **(b)** Frequency of co-mutational patterns in *NPM1*$^{\text{class I}}$ for the five most commonly mutated genes in NPM1 overall. **(c)** Frequency of co-mutational patterns in *NPM1*$^{\text{class II}}$ for the four most commonly mutated genes in NPM1 overall.

(a)

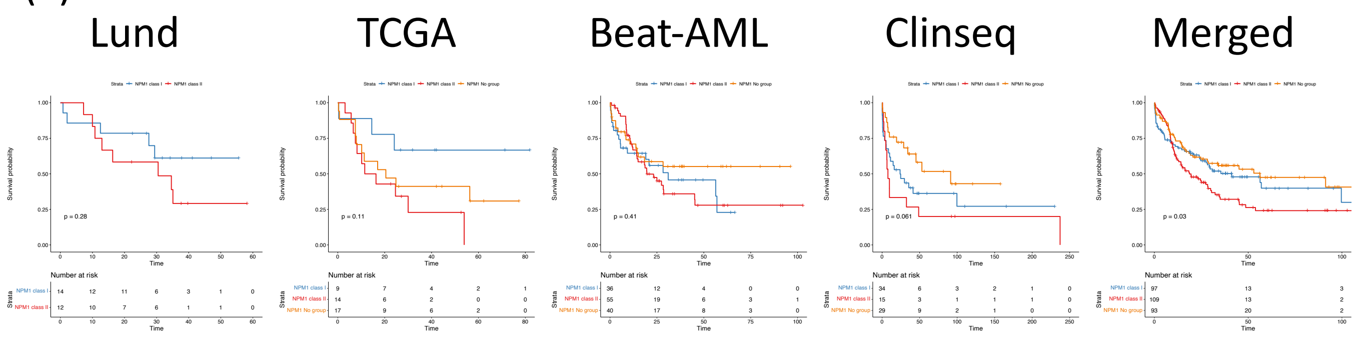


(b)

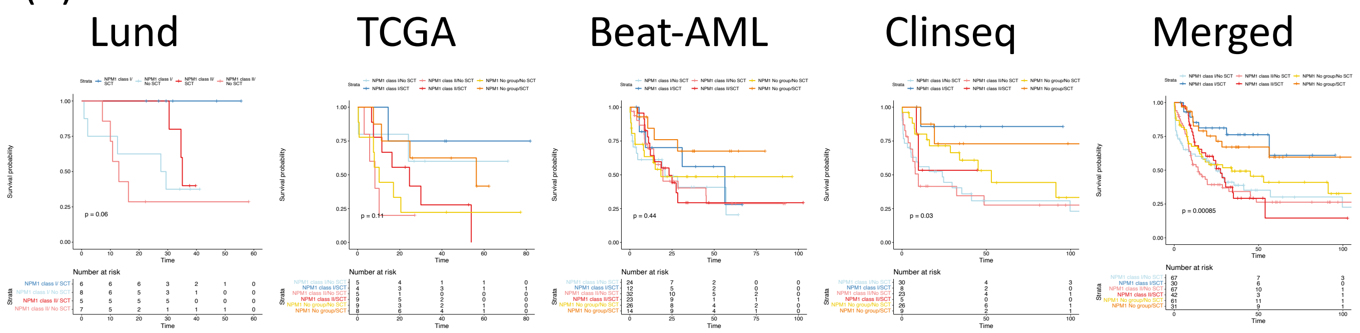


(c)

| Variable | | N | Hazard ratio | | p |
|---|---|---|---|---|---|
| **NPM1group** | NPM1 class I | 79 | | Reference | |
| | NPM1 class II | 95 | | 1.87 (1.20, 2.92) | 0.006 |
| | NPM1 No group | 76 | | 1.30 (0.78, 2.18) | 0.314 |
| **Age.group** | high | 135 | | Reference | |
| | low | 115 | | 0.84 (0.58, 1.22) | 0.354 |
| **WBC.group** | high | 111 | | Reference | |
| | low | 139 | | 0.87 (0.60, 1.25) | 0.450 |
| **SCT.group** | no | 154 | | Reference | |
| | yes | 96 | | 0.39 (0.26, 0.59) | <0.001 |
| **FLT3.ITD.group** | neg | 128 | | Reference | |
| | pos | 122 | | 2.87 (1.92, 4.29) | <0.001 |

**Supplementary Fig. 14 | Overall survival for *NPM1*$^{class\ I}$ and *NPM1*$^{class\ II}$ subtypes. (a)** Overall survival for all patients with *NPM1* mutated AML that were treated with curative intent in the Lund, TCGA, Beat-AML, and Clinseq studies shown as Kaplan-Meier curves for each dataset separately (left) and merged together (right). **(b)** Overall survival for all patients with *NPM1* mutated AML that were treated with curative intent in the Lund, TCGA, Beat-AML, and Clinseq studies, as in (a), but with separate curves for patients that received and did not receive a hematopoietic stem cell transplantation. **(c)** Forest plot illustrating the hazard ratio from a multivariate Cox proportional hazard ratio of the overall survival. The following covariates are included NPM1group (*NPM1*$^{class\ I}$, *NPM1*$^{class\ II}$, *NPM1* No group), Age.group (high: >= 60 years, low: < 60 years), WBC.group (high: >42x10$^9$/L; low: <= 42x10$^9$/L), SCT.group (no: no stem cell transplantation; yes: received stem cell transplantation), and FLT3.ITD.group (neg: negative for *FLT3*-ITD; pos: positive for *FLT3*-ITD).

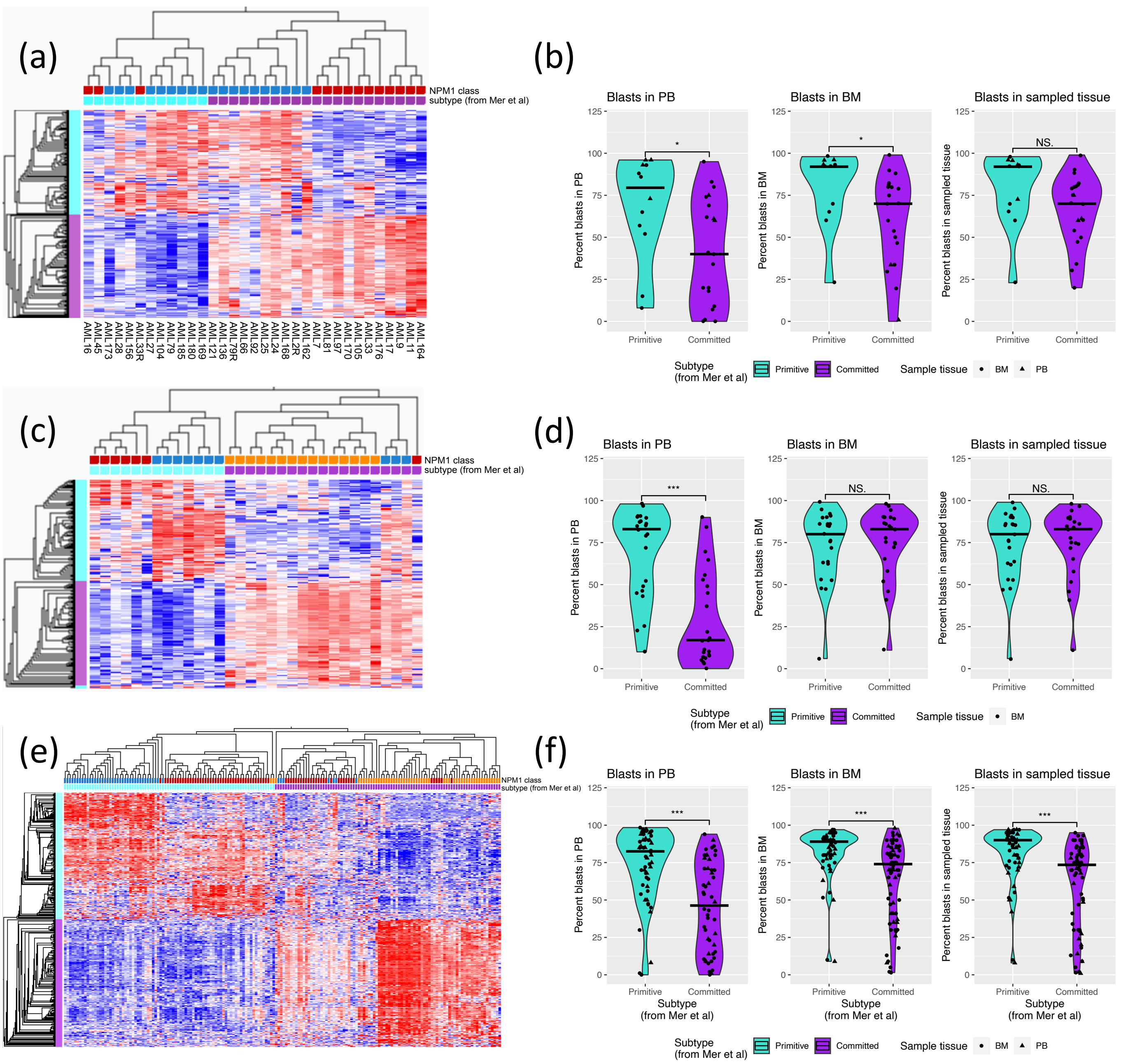


**Supplementary Fig. 15 | Primitive and committed subtypes described by Mer et. al does not overlap with *NPM1*$^{\text{class I}}$ and *NPM1*$^{\text{class II}}$ subtypes. (a)** Hierarchical clustering of bulk gene expression data from 33 *NPM1*-mutated AML samples from the Lund dataset based on the expression of the 100 most highly expressed genes in the primitive subtype (genes indicated in turquoise) and the 100 most highly expressed genes in the committed subtype (genes indicated in purple), as described by Mer et al.[7] *NPM1*$^{\text{class I}}$ (blue)/*NPM1*$^{\text{class II}}$ (red) classification and primitive (turquoise)/committed (purple) classification (based on this hierarchical clustering) is indicated above the heatmap. **(b)** Percent blasts in PB, BM, and sample tissue (either PB or BM) for primitive and committed cases from Lund, with significant differences indicated by asterisks (Mann-Whitney U test). **(c)** Hierarchical clustering of bulk gene expression data from 48 *NPM1*-mutated AML samples from the TCGA dataset[1] based on the expression of the 100 most highly expressed genes in the primitive subtype (genes indicated in turquoise) and the 100 most highly expressed genes in the committed subtype (genes indicated in purple), as described by Mer et al.[7] *NPM1*$^{\text{class I}}$ (blue)/*NPM1*$^{\text{class II}}$ (red) classification and primitive (turquoise)/committed (purple) classification (based on this hierarchical clustering) is indicated above the heatmap. **(d)** Percent blasts in PB, BM, and sample tissue (either PB or BM) for primitive and committed cases from TCGA, with significant differences indicated by asterisks (Mann-Whitney U test). **(e)** Hierarchical clustering of bulk gene expression data from 174 *NPM1*-mutated AML samples from the Beat-AML 2.0 dataset[5] based on the expression of the 300 most highly expressed genes in the primitive subtype (genes indicated in turquoise) and the 300 most highly expressed genes in the committed subtype (genes indicated in purple), specifically in the Beat-AML cohort as described by Mer et al.[7] *NPM1*$^{\text{class I}}$ (blue)/*NPM1*$^{\text{class II}}$ (red) classification and primitive (turquoise)/committed (purple) classification (based on this hierarchical clustering) is indicated above the heatmap. **(f)** Percent blasts in PB, BM, and sample tissue (either PB or BM) for primitive and committed cases from Beat-AML 2.0, with significant differences indicated by asterisks (Mann-Whitney U test).

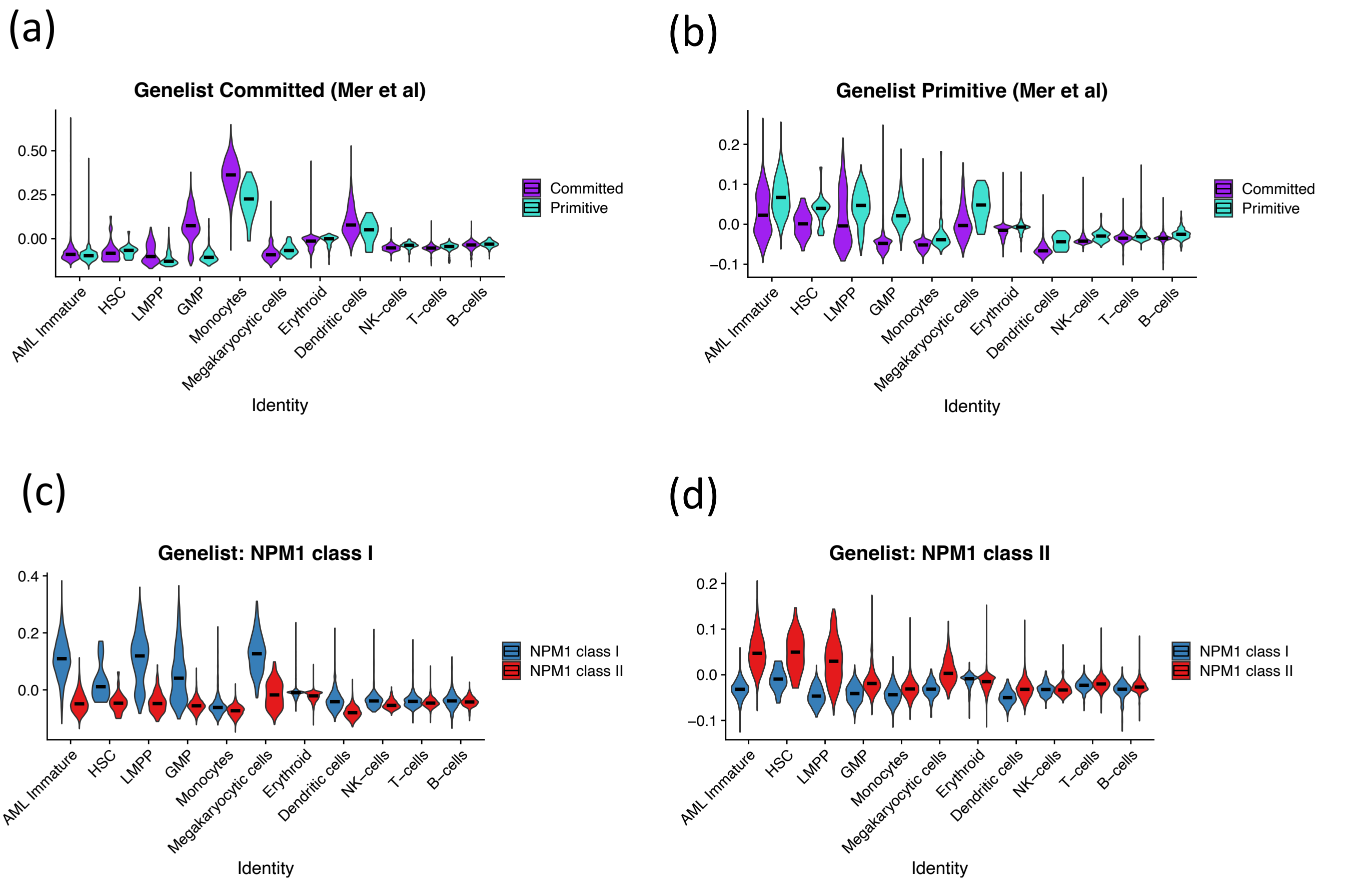


**Supplementary Fig. 16 | Gene expression across cell types for genes associated with *NPM1* subtypes. (a)** Average expression of genes defining the committed subtype across cell types in single cell data from twelve *NPM1*-mutated AML samples, divided into committed and primitive samples. The gene expression module is defined by the top 100 overexpressed genes in the committed subtype, as described by Mer et al.[7] **(b)** Average expression of genes defining the primitive subtype across cell types in single cell data from twelve *NPM1*-mutated AML samples, divided into committed and primitive samples. The gene expression module is defined by the top 100 overexpressed genes in the primitive subtype, as described by Mer et al.[7] **(c)** Average expression of genes defining *NPM1*$^{\text{class I}}$ across cell types in single cell data from *NPM1*-mutated AML samples, divided into *NPM1*$^{\text{class I}}$ and *NPM1*$^{\text{class II}}$. The gene expression module is defined by 79 genes specifically expressed in immature cells in *NPM1*$^{\text{class I}}$ samples. **(d)** Average expression of genes defining *NPM1*$^{\text{class II}}$ across cell types in single cell data from *NPM1*-mutated AML samples, divided into *NPM1*$^{\text{class I}}$ and *NPM1*$^{\text{class II}}$. The gene expression module is defined by 101 genes specifically expressed in immature cells in *NPM1*$^{\text{class II}}$ samples.

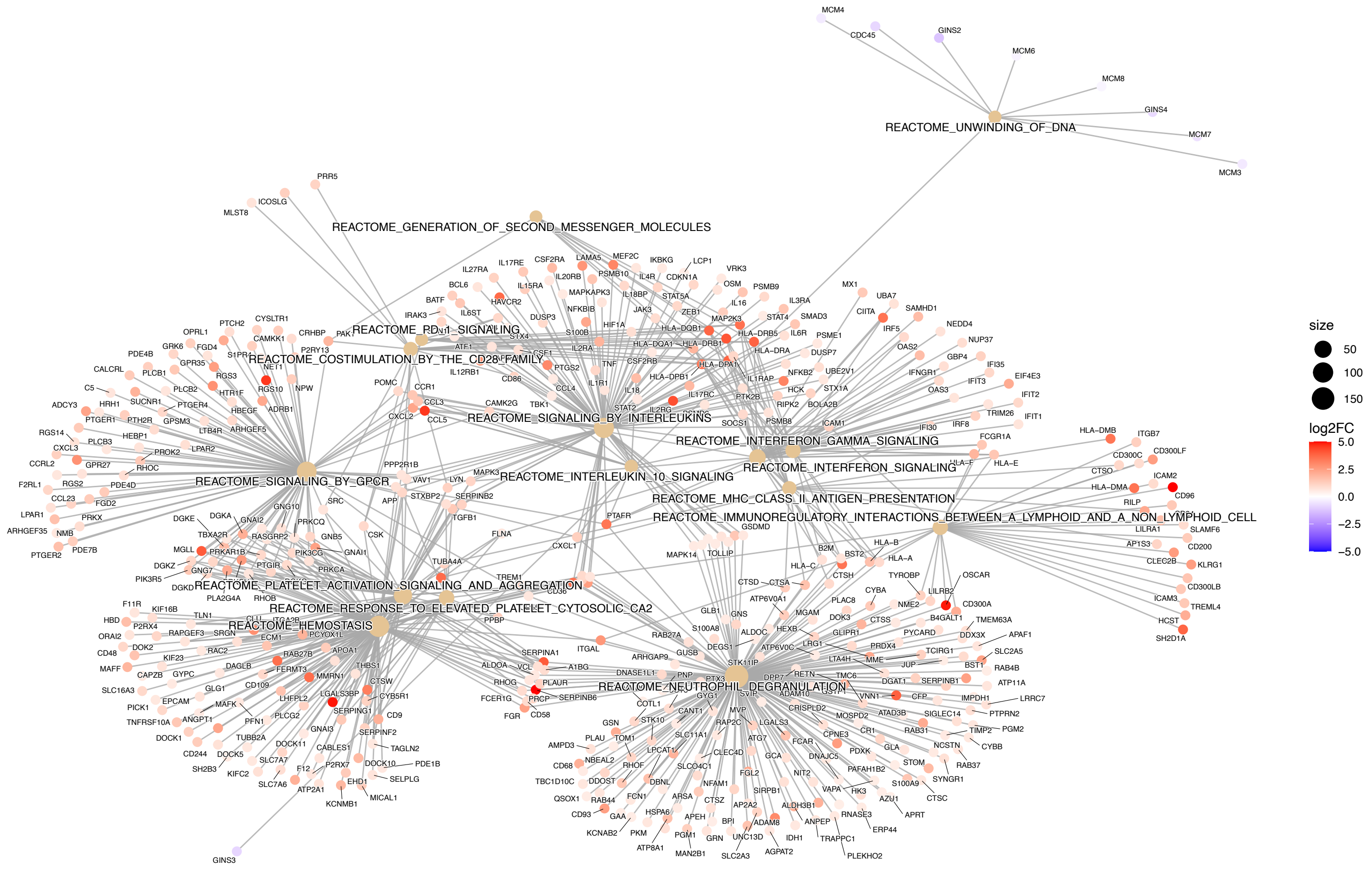


**Supplementary Fig. 17 | Gene set enrichment analysis between *NPM1*$^{\text{class I}}$ and *NPM1*$^{\text{class II}}$ subtypes.** Network plot visualizing gene set enrichment analysis for differentially expressed genes between AML immature cells in *NPM1*$^{\text{class I}}$ and *NPM1*$^{\text{class II}}$ samples. The fifteen most enriched of the msigdb canonical pathways reactome gene sets are included, together with core enriched genes from these gene sets.

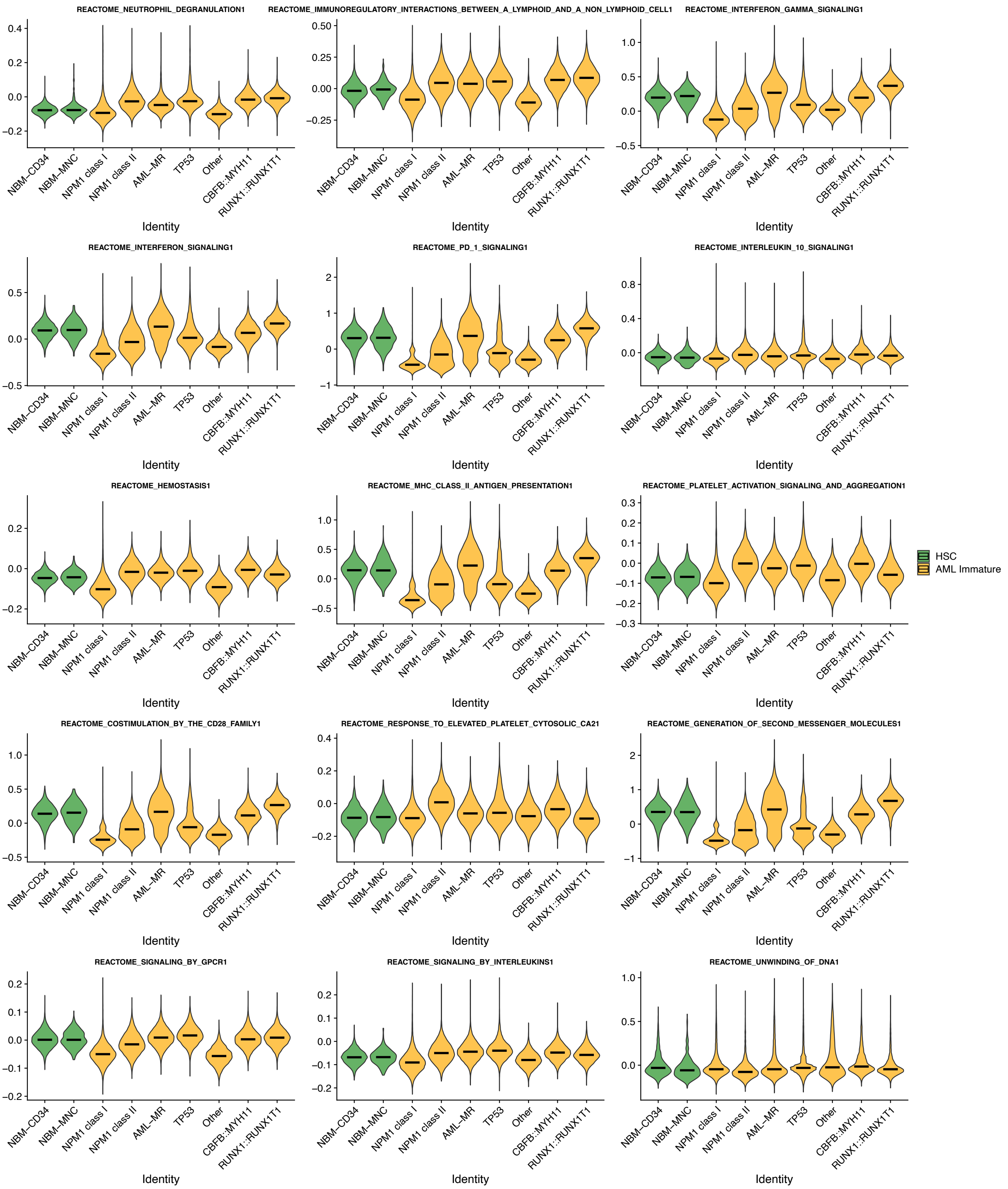


**Supplementary Fig. 18 | Expression level of core expressed genes from GSEA enriched pathways in AML immature cells and NBM HSC cells.** Expression level of modules defined by the core enriched genes from the fifteen most enriched gene sets from reactome is visualized in HSC cells from NBM-CD34 and NBM-MNC samples and in AML immature cells from different AML subtypes.

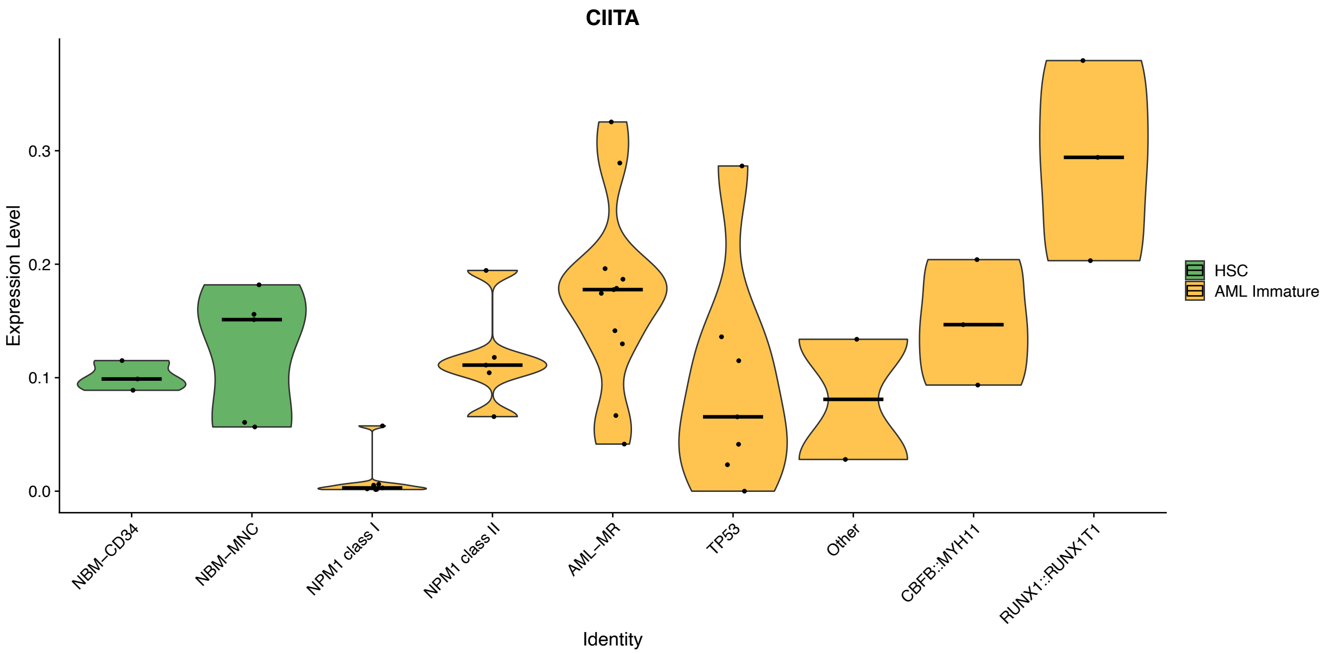


**Supplementary Fig. 19 | Average expression of CIITA in AML immature cells and NBM HSC cells.** The average expression of CIITA in AML immature cells from different AML subtypes and HSC cells from NBM-CD34 and NBM-MNC samples.

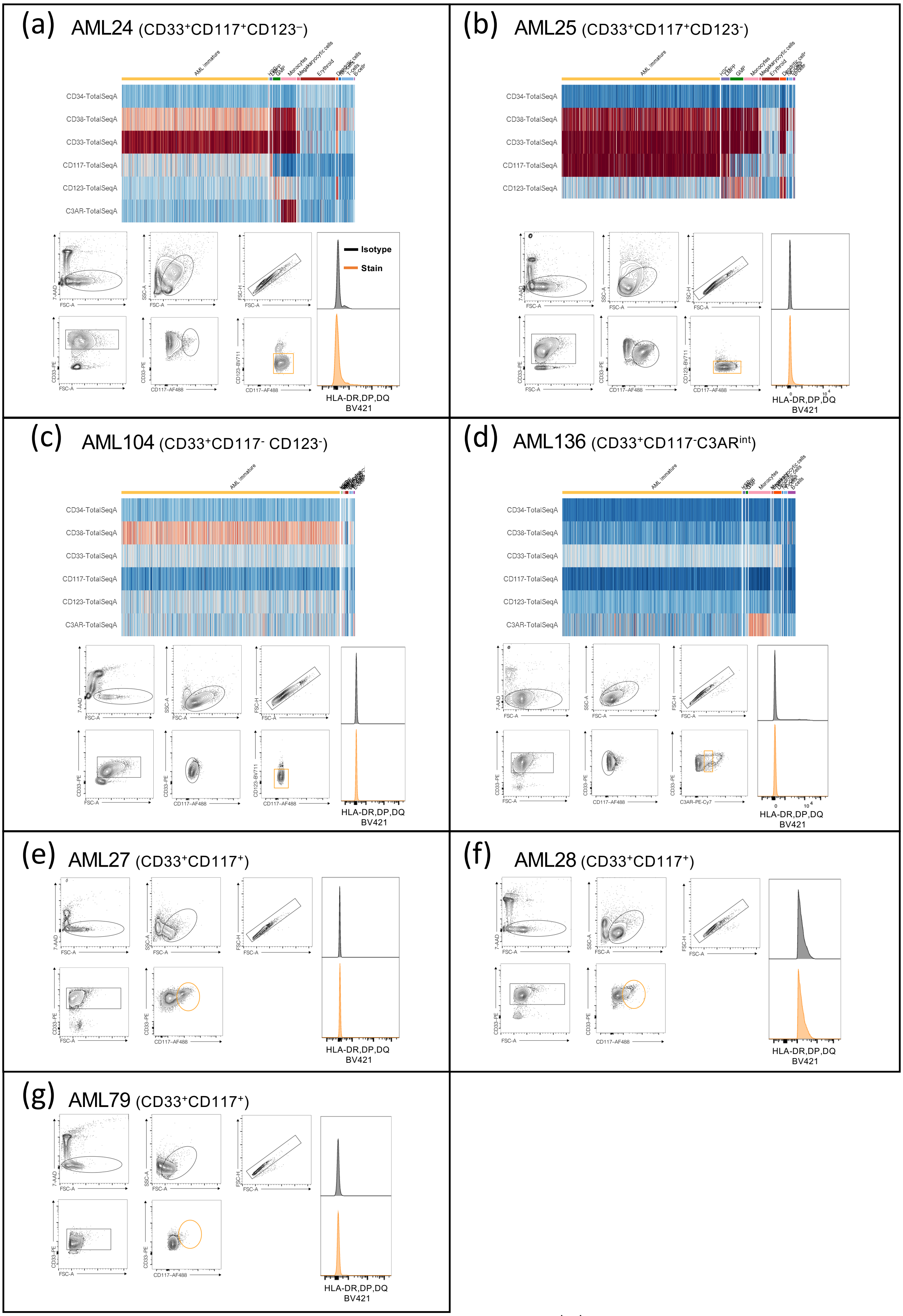


**Supplementary Fig. 20 | Cell surface markers used to define AML immature cells in *NPM1*$^{class\ I}$ samples. (a-d)** Heatmaps illustrating the expression of cell surface markers in AML24, AML25, AML104 and AML136 as identified by scADT-seq (top). Scatter plots showing the identification of AML immature cells by FACS (bottom left). Histograms showing HLA-DR, -DP, and -DQ expression in the AML immature cells (bottom right). **(e-g)** Scatter plots showing the identification of AML immature cells in AML27, AML28 and AML79 by FACS, defined as $CD33^+CD117^+$ (left). Histograms showing HLA-DR, -DP, and -DQ expression in the AML immature cells of AML27, AML28 and AML79 (right).

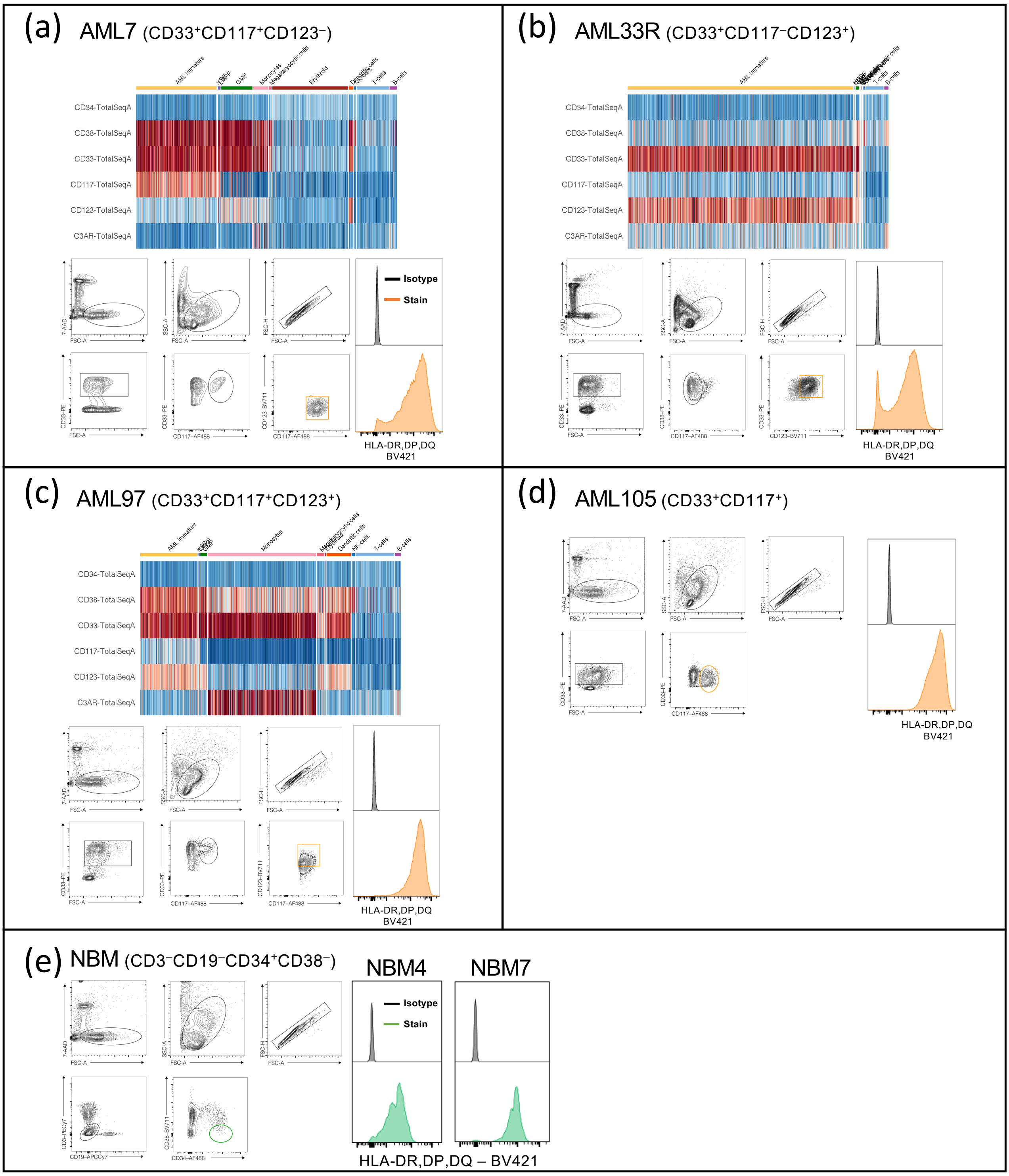


**Supplementary Fig. 21 | Cell surface markers used to define AML immature cells in *NPM1*$^{\text{class II}}$ samples. (a-c)** Heatmaps illustrating the expression of selected cell surface markers in AML7, AML33R, and AML97 as identified by scADT-seq (top). Scatter plots showing the identification of AML immature cells by FACS using immune profiles defined by scADT-seq (bottom left). Histograms showing HLA-DR, -DP, and -DQ expression in the AML immature cells (bottom right). **(d)** Scatter plots showing the identification of AML immature cells in AML105, defined as CD33$^+$CD117$^+$, by FACS (left). Histograms showing HLA-DR, -DP, and -DQ expression in the AML immature cells of AML105 (right). **(e)** Scatter plots showing the identification of immature (CD34$^+$CD38$^-$) cells from NBM using FACS. Histograms showing HLA-DR, -DP, and -DQ expression in CD34$^+$CD38$^-$ cells from two NBM samples.

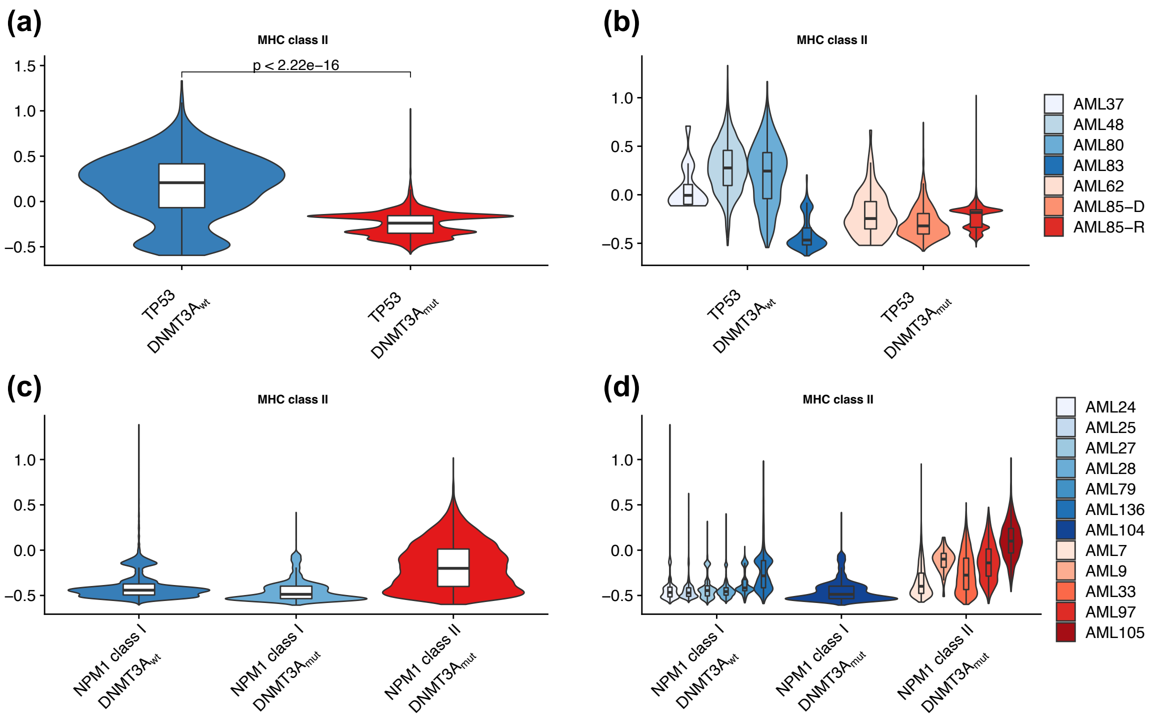


**Supplementary Fig. 22 | Expression of genes encoding MHC class II components in AML immature cells from *TP53*-mutated and *NPM1*$^{\text{class I}}$/*NPM1*$^{\text{class II}}$ samples divided by *DNMT3A* mutation status. (a-b)** Single cell expression of genes encoding MHC class II components in AML immature cells from *TP53*-mutated AMLs harboring wildtype *DNMT3A* (*TP53*/$DNMT3A_{wt}$, 4 cases) compared with mutated DNMT3A (*TP53*/$DNMT3A_{mut}$, 3 cases). The data is presented with all samples grouped together (a) and individually for each sample (b). The expression of genes encoding MHC class II components in this subtype is significantly higher in *DNMT3A* wild type cases. **(c-d)** Single cell expression of genes encoding MHC class II components compared between AML immature cells from *NPM1*$^{\text{class I}}$/*DNMT3A* wildtype (6 cases), *NPM1*$^{\text{class I}}$/*DNMT3A* mutated (1 case), and *NPM1*$^{\text{class II}}$/*DNMT3A* mutated AMLs (5 cases). The data is presented with all samples grouped together (c) and individually for each sample (d). The data set did not include any *NPM1*$^{\text{class II}}$/*DNMT3A* wildtype cases. The single *NPM1*$^{\text{class I}}$/*DNMT3A* mutated case exhibits expression of MHC class II components on the same level or below other *NPM1*$^{\text{class I}}$ AMLs, and clearly below the level of *NPM1*$^{\text{class II}}$ AMLs. The data from both *TP53*-mutated and *NPM1*-mutated AMLs indicate that high MHC class II expression (as observed in *NPM1*$^{\text{class II}}$ AMLs) is not a general feature of AML with mutated *DNMT3A*.

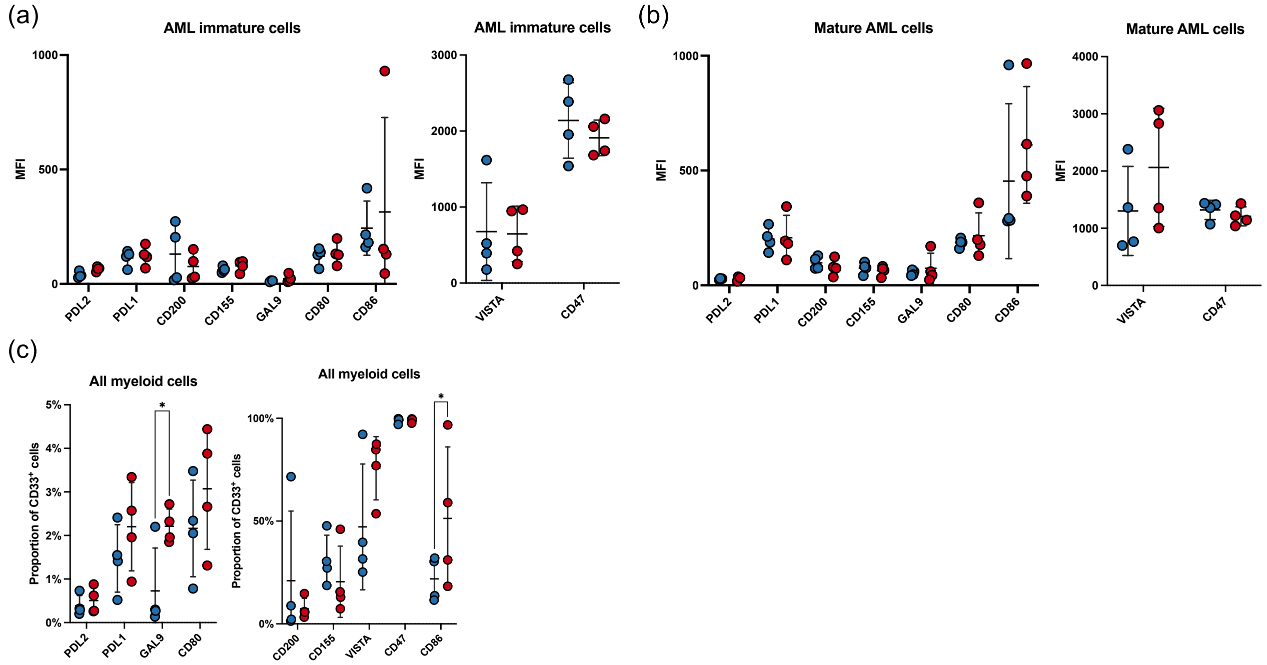


**Supplementary Fig. 23 | Checkpoint receptor surface expression on *NPM1*$^{\text{class I}}$ and *NPM1*$^{\text{class II}}$ AML cells. (a)** Surface expression of selected checkpoint receptors on immature (CD123+) AML cells from *NPM1*$^{\text{class I}}$ (blue) and *NPM1*$^{\text{class II}}$ (red) measured by flow cytometry. **(b)** Surface expression of selected checkpoint receptors on mature (CD14+) AML cells from *NPM1*$^{\text{class I}}$ (blue) and *NPM1*$^{\text{class II}}$ measured by flow cytometry. **(c)** Proportion of myeloid (CD33+) AML cells positive for selected checkpoint receptors as measured by flow cytometry for *NPM1*$^{\text{class I}}$ (blue) and *NPM1*$^{\text{class II}}$ (red).

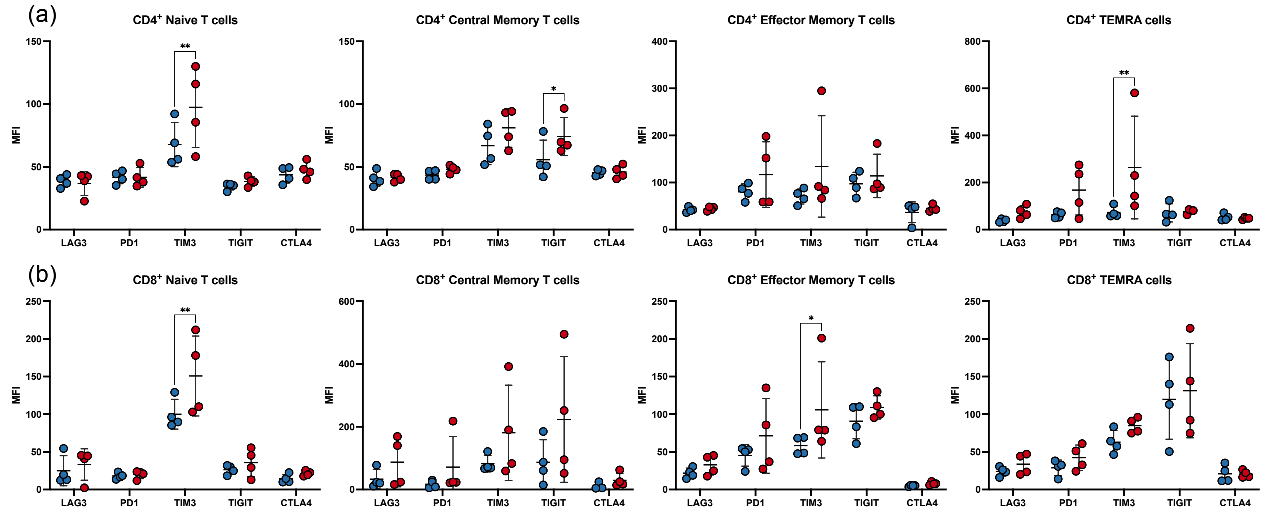


**Supplementary Fig. 24 | Checkpoint receptor surface expression on T cell subsets from *NPM1*$^{class\ I}$ and *NPM1*$^{class\ II}$ diagnostic bone marrow. (a)** Surface expression of selected checkpoint molecules on $CD4^+$ naive T cells ($CCR7^+CD45RA^+$; first panel), $CD4^+$ central memory T cells ($CCR7^+CD45RA^-$; second panel), $CD4^+$ effector memory T cells ($CCR7^-CD45RA^-$; third panel), and $CD4^+$ T effector memory cells re-expressing CD45RA (TEMRA cells; $CCR7^-CD45RA^+$; fourth panel) from *NPM1*$^{class\ I}$ (blue) and *NPM1*$^{class\ II}$ (red) diagnostic bone marrow, measured by flow cytometry. **(b)** Surface expression of selected checkpoint molecules on $CD8^+$ naive T cells ($CCR7^+CD45RA^+$; first panel), $CD8^+$ central memory T cells ($CCR7^+CD45RA^-$; second panel), $CD8^+$ effector memory T cells ($CCR7^-CD45RA^-$; third panel), and $CD8^+$ T effector memory cells re-expressing CD45RA (TEMRA cells; $CCR7^-CD45RA^+$; fourth panel) from *NPM1*$^{class\ I}$ (blue) and *NPM1*$^{class\ II}$ (red) diagnostic bone marrow, measured by flow cytometry.

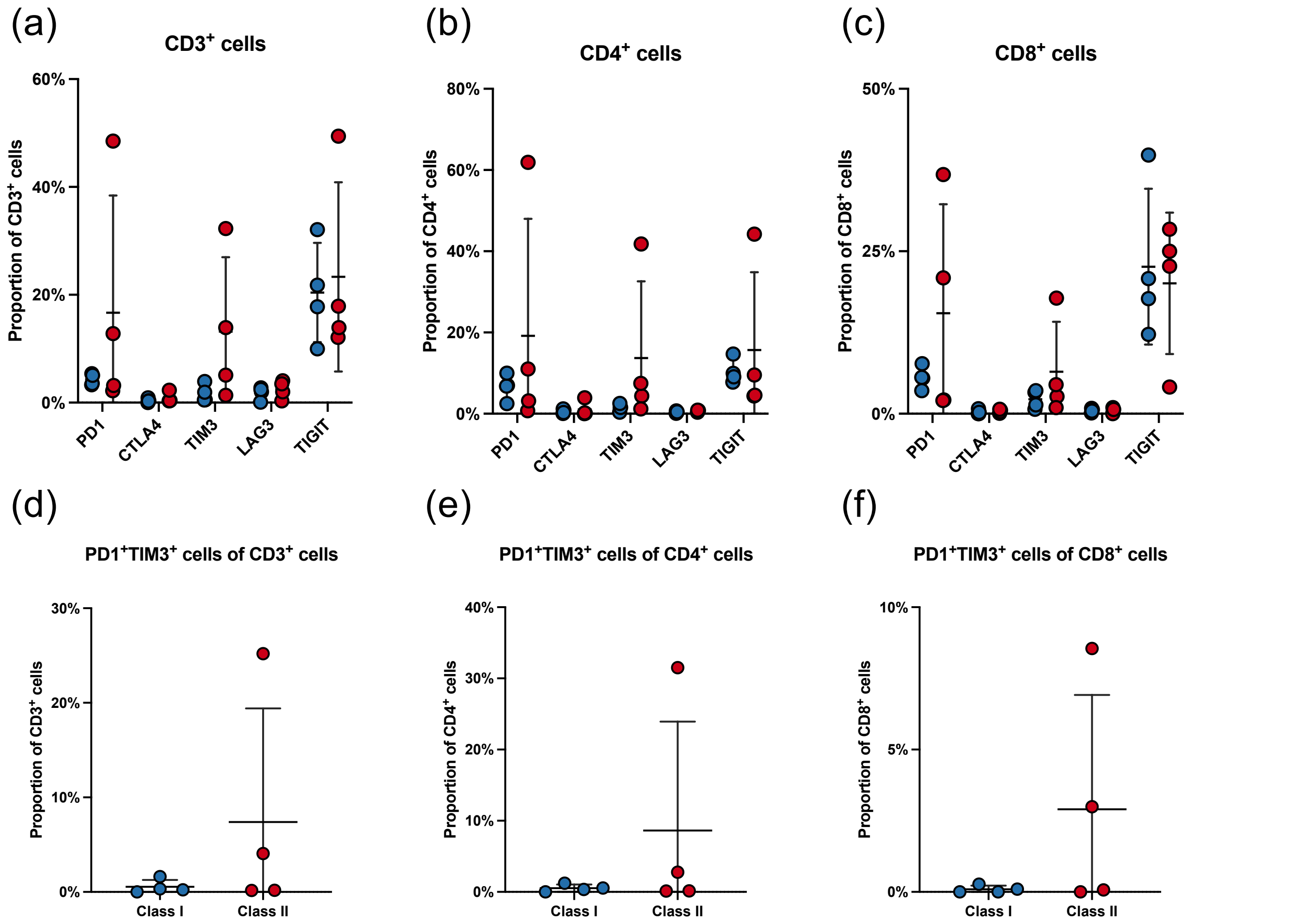


**Supplementary Fig. 25 | Proportion of T cells expressing checkpoint receptor combinations in *NPM1*$^{\text{class I}}$ and *NPM1*$^{\text{class II}}$ diagnostic bone marrow. (a)** Proportion of T cells (CD3+) from AML diagnostic bone marrow positive for selected checkpoint receptors, as measured by flow cytometry for *NPM1*$^{\text{class I}}$ (blue) and *NPM1*$^{\text{class II}}$ (red). **(b)** Proportion of CD4$^+$ T cells from AML diagnostic bone marrow positive for selected checkpoint receptors, as measured by flow cytometry for *NPM1*$^{\text{class I}}$ (blue) and *NPM1*$^{\text{class II}}$ (red). **(c)** Proportion of CD8$^+$ T cells from AML diagnostic bone marrow positive for selected checkpoint receptors, as measured by flow cytometry for *NPM1*$^{\text{class I}}$ (blue) and *NPM1*$^{\text{class II}}$ (red). **(d)** Proportion of T cells (CD3+) from AML diagnostic bone marrow expressing both PD1 and TIM3, indicating an exhausted phenotype, as measured by flow cytometry for *NPM1*$^{\text{class I}}$ (blue) and *NPM1*$^{\text{class II}}$ (red). **(e)** Proportion of CD4$^+$ T cells from AML diagnostic bone marrow expressing both PD1 and TIM3, indicating an exhausted phenotype, as measured by flow cytometry for *NPM1*$^{\text{class I}}$ (blue) and *NPM1*$^{\text{class II}}$ (red). **(f)** Proportion of CD8$^+$ T cells from AML diagnostic bone marrow expressing both PD1 and TIM3, indicating an exhausted phenotype, as measured by flow cytometry for *NPM1*$^{\text{class I}}$ (blue) and *NPM1*$^{\text{class II}}$ (red).

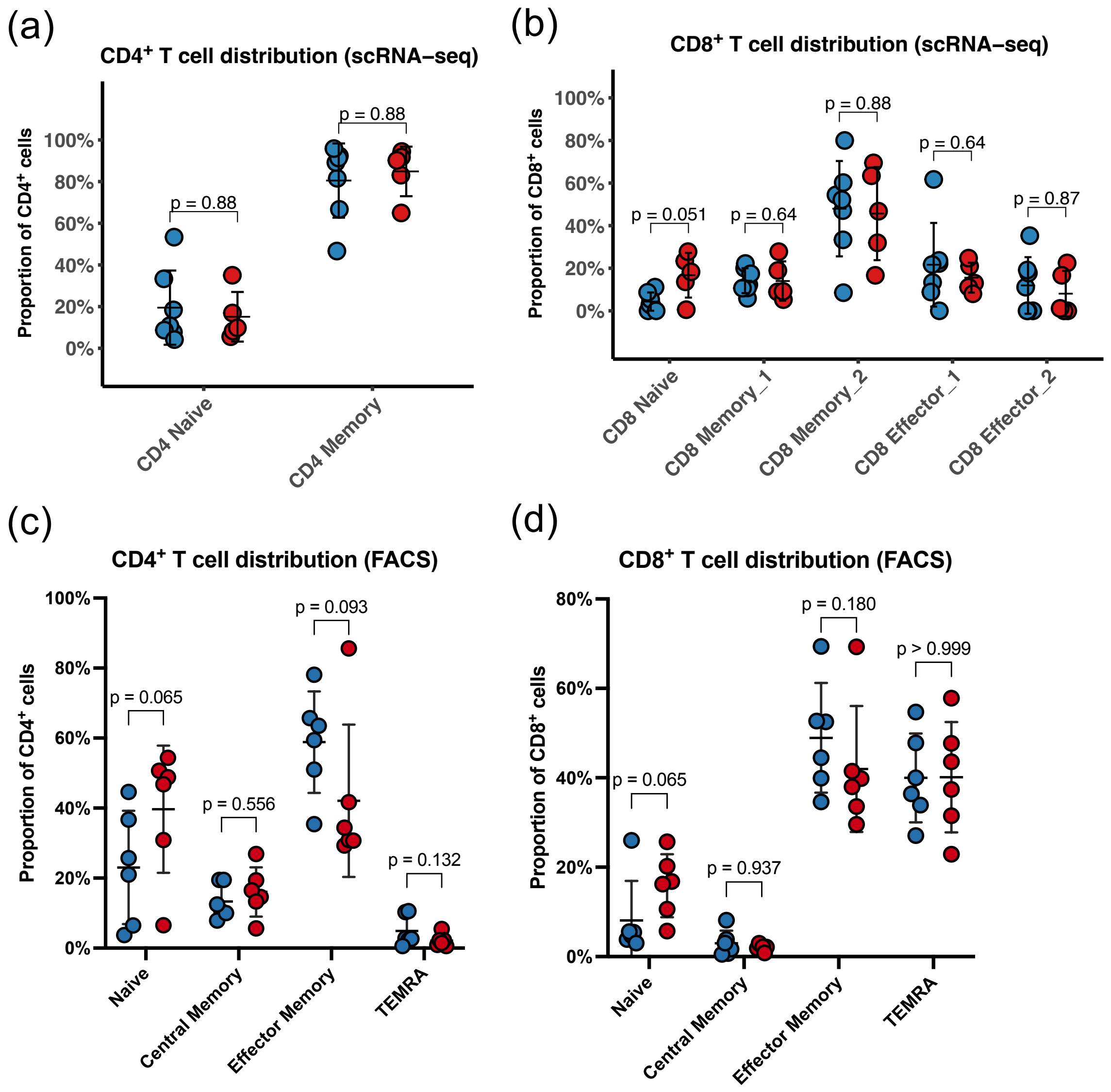


**Supplementary Fig. 26 | Proportion of T cell subsets in *NPM1*$^{\text{class I}}$ and *NPM1*$^{\text{class II}}$ samples.** **(a)** Composition of the CD4$^+$ T cell compartment in *NPM1*$^{\text{class I}}$ (blue) and *NPM1*$^{\text{class II}}$ (red) BM or PB as determined by scRNA-seq. **(b)** Composition of the CD8$^+$ T cell compartment in *NPM1*$^{\text{class I}}$ (blue) and *NPM1*$^{\text{class II}}$ (red) BM or PB as determined by scRNA-seq. **(c)** Composition of the CD4$^+$ T cell compartment in *NPM1*$^{\text{class I}}$ (blue) and *NPM1*$^{\text{class II}}$ (red) BM as determined by flow cytometry. **(d)** Composition of the CD8$^+$ T cell compartment in *NPM1*$^{\text{class I}}$ (blue) and *NPM1*$^{\text{class II}}$ (red) BM as determined by flow cytometry.
